# Lineage hierarchies and stochasticity ensure the long-term maintenance of adult neural stem cells

**DOI:** 10.1101/663922

**Authors:** Emmanuel Than-Trong, Bahareh Kiani, Nicolas Dray, Sara Ortica, Benjamin Simons, Steffen Rulands, Alessandro Alunni, Laure Bally-Cuif

## Abstract

The cellular basis and extent of neural stem cell (NSC) self-renewal in adult vertebrates, and their heterogeneity, remain controversial. To explore the functional behavior and dynamics of individual NSCs within brain germinal pools, we combined genetic lineage tracing, quantitative clonal analysis, intravital imaging and global population assessments in the adult zebrafish telencephalon. We show that adult neurogenesis is organized in a hierarchy where a subpopulation of reservoir NSCs with longterm self-renewal potential generate a pool of operational NSCs taking stochastic fates biased towards neuronal differentiation. To fuel the long-term growth of the adult germinal niche, we provide evidence for the existence of an additional, upstream, progenitor population that supports the continuous generation of new reservoir NSCs, contributing to their overall expansion. Hence, the dynamics of vertebrate neurogenesis relies on a hierarchical organization where growth, self-renewal and neurogenic functions are segregated between different NSC types.

## Report

The brain of most adult vertebrate species, including human, hosts specialized precursor cells, called neural stem cells (NSCs), which fuel the ongoing production of neurons into discrete brain regions (1–5). Adult-born neurons bring an additional layer of plasticity into local circuits that appears critical for particular aspects of learning and memory (6). In mammals, alterations of adult neurogenesis have been linked functionally to major depression and anxiety-like behaviors (6, 7). Despite extensive studies, the functional and molecular identity, long-term renewal potential, and pattern of division of NSCs remain controversial. Notably, while several works suggest that NSCs in both the dentate gyrus (DG) and the subependymal zone (SEZ) of the mammalian telencephalon are progressively consumed over time (8–13), others report a substantial self-renewal (14–16), or even amplification, potential of NSCs (17). Similar discrepancies were also reported in the adult zebrafish telencephalon (18, 19). Accordingly, most models put forward in these studies also reached conflicting conclusions regarding the mode of NSC divisions. Whether such divergences reflect technical specificities and/or represent an underlying heterogeneity of the NSC compartment remains an important open question.

To resolve the functional identity, fate behavior and lineage dependencies of NSCs in an adult vertebrate brain, we used quantitative clonal analysis to target their fate in the zebrafish dorsal telencephalon (pallium). The everted morphology of the pallium exposes its ventricle (Fig. S1A), making constituent NSCs and their progeny readily accessible to whole-mount investigation, and allowing a combination of intravital imaging and genetic cell lineage tracing over a lifetime (20, 21). Further, under physiological conditions, progenitor cell migration is never observed and newly-generated neurons delaminate to settle under the pallial ventricular surface adjacent to the place where they were born (19, 22). Thus, the zebrafish pallium provides an optimal system to perform clonal analysis of NSC fates, as neural clones remain compact and superficial, allowing their whole cellular complement to be captured in a single *in toto* confocal acquisition.

Zebrafish pallial NSCs share the same basic regulatory mechanisms and physiological requirements as mammalian NSCs (23). Based on markers, these astroglial cells are thought to be characterized by the expression of the glial fibrillary acidic protein (Gfap) and the enzyme glutamine synthetase (Gs) -whose expression is redundant with Gfap-, the Notch-target gene *hairy-related 4 (her4.1* – orthologous to mammalian *Hes5*), and the stem cell-associated transcription factor Sox2 (5, 20, 24–27). *her4.1*:dRFP and *gfap.nGFP* expression in the pallium of double transgenic fish almost completely overlap and appear to characterize the same, mostly quiescent, NSC population, which accounts for about 75% of Sox2-positive progenitors (Fig. 1A and S1A-D). The remaining Sox2+ cells express neither *her4.1* nor *gfap* and are, for the majority (57.7±2.1%), in an activated/proliferating state (identifying them as activated non-astroglial neural progenitors -aNPs-, against 42.3±2.1% quiescent/non-proliferating neural progenitors -qNPs) (Fig. S1D). aNPs make up the bulk of actively proliferating pallial progenitors (Fig. S1A,E), likely representing transit amplifying progenitors (TAPs), which generate neurons after a limited number of cell divisions (19, 25, 26).

**Fig. 1.**
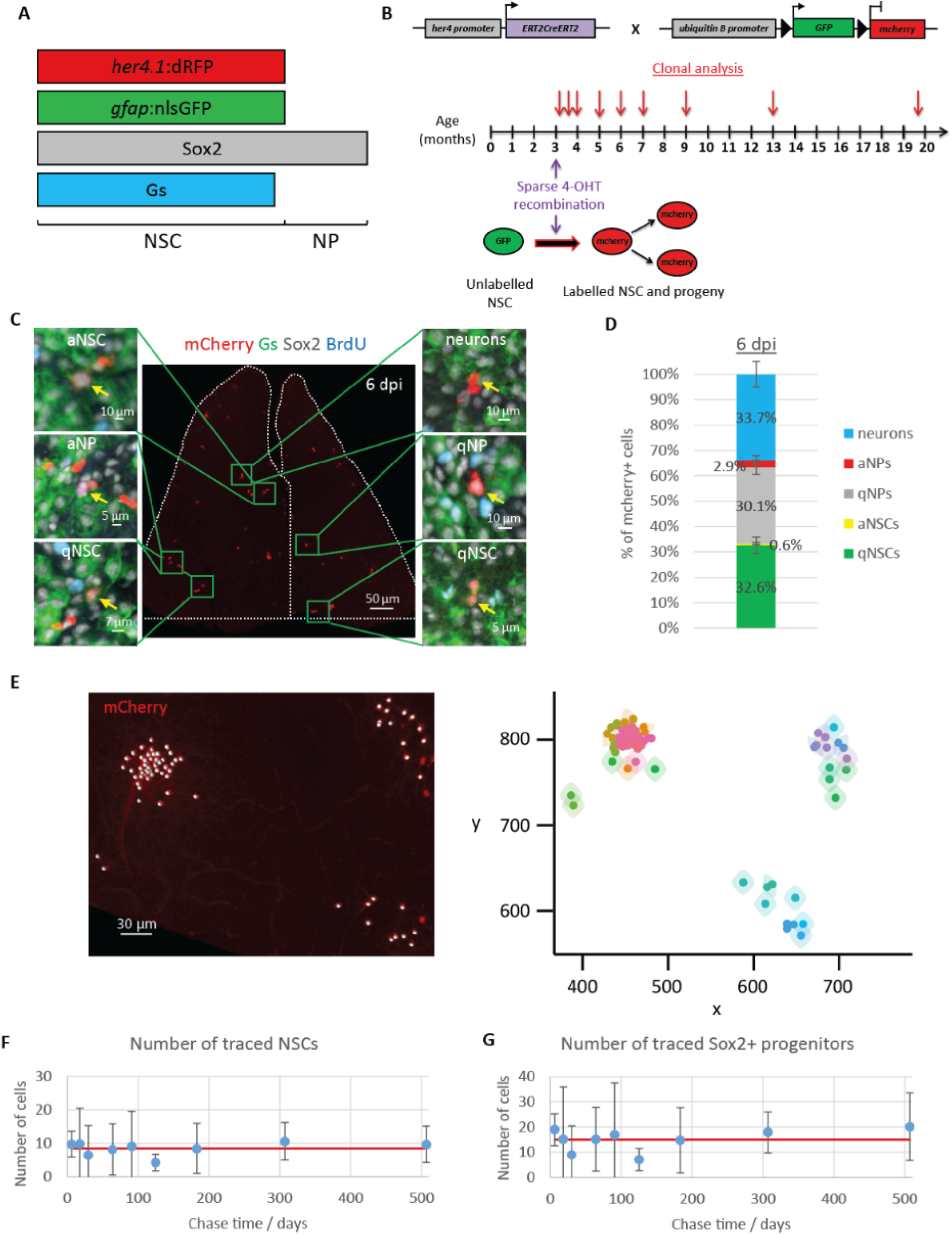
Lineage tracing of *her4.1-expressing* NSCs shows homeostasis. **(A)** Summary of markers characterizing Dm pallial progenitors. **(B)** Cre recombination was sparsely induced in 3 mpf (months post-fertilization) *her4.1:iCre;ubi:Switch* double transgenic adults by 4-hydroxytamoxifen (4-OHT), resulting in mCherry expression in recombined NSCs and their progeny. Analysed time points (arrows) span between 6 and 507 days-post-induction (dpi). **(C)** Dorsal view of a representative pallium showing sparsely induced cells at 6 dpi (dotted area to the pallial Dm territory of interest, Fig. S5). Boxed areas are magnified to illustrate the different cell types traced (yellow arrows). Proliferating progenitors were labelled by a 24h BrdU pulse. NSCs are Gs^+^ and Sox2+, NPs are Sox2^+^ only. In contrast to aNSCs and aNPs, qNSCs and qNPs did not incorporate BrdU during the pulse. **(D)** Cell type composition of the mCherry^+^ population at 6 dpi. n=6 brains. **(E)** Example of clustering at 183 dpi. Left: dorsal view of a three-dimensional reconstruction of a hemisphere. The spots registering the coordinates of traced cells (red) are shown (white dots). Right: two-dimensional map of the clones identified by the clustering algorithm. Clones are color-coded. **(F)** Average number of traced (mCherry^+^) Gs+, Sox2^+^ NSCs across all time points analyzed. One-way ANOVA: F_(8,37)_=1, p=0.45; all pairwise comparisons: LSD test followed by Holm’s adjustment. Error bars: 95% CI. **(G)** Average number of traced (mCherry^+^) Sox2^+^ progenitors across all time points analyzed. One-way ANOVA: F_(8,37)_=1.65, p=0.14; all pairwise comparisons: LSD test followed by Holm’s adjustment. Error bars: 95% CI. **(F, G)** n= 6, 3, 3, 3, 5, 6, 7, 8 and 6 brains at 6, 18, 30, 64, 91, 125, 183, 307 and 507 dpi, respectively.

To target the fate of individual pallial NSCs and their progeny in the dorso-medial pallial domain (Dm) during adulthood over extended periods of time, we opted for an inducible genetic lineage tracing approach (Fig. 1B). We crossed the transgenic driver line *Tg(her4.1:ERT2CreERT2)* (for short: *her4.1:iCre)*, which expresses a tamoxifen-inducible Cre recombinase in *her4.1* -expressing NSCs (Fig. S2), with the *Tg(−3.5ubb:loxP-EGFP-loxP-mCherry)* (for short: *ubi:Switch)* reporter line (28) (Fig. S3). To establish the conditions for clonal induction, we induced 3-month post-fertilization (mpf) adult zebrafish with decreasing concentrations and exposure times to 4-hydroxytamoxifen (4-OHT) until reaching an average number of 20.7±2.45 (mean ± s.e.m.) cell clusters per hemisphere at 6 days postinduction (dpi) (Fig. 1B-D, S4, S5 and supplementary text). Under these conditions, by 6 dpi, only 1.6% ± 0.3% of the total Sox2+ cell population was labelled, while 46% of clones had already divided and/or differentiated into neurons (Fig. S6). To assign the clonal provenance of labelled cells with known statistical confidence, we combined a hierarchical clustering approach with a biophysical model of clone dispersion (Fig. 1E, S7 and supplementary text).

Stem cells are defined functionally by their long-term self-renewal capacity. Following the report that zebrafish pallial NSCs might be progressively consumed through direct differentiation into neurons (18), in apparent contradiction with their measured steady density over the considered time frame (29), we first explored their maintenance within the *her4* lineage. Notably, we found that the total number of *her4.1:iCre-traced* Gs^+^ Sox2^+^ NSCs, averaged across animals, was maintained at a seemingly constant level throughout the ~ 17-month chase period spanning more than half the adult zebrafish life (Fig. 1F). This result also applied to the whole population of traced Sox2^+^ progenitors (Fig. 1G). Thus, the targeted *her4* lineage is, as a whole, homeostatic and self-reliant for its maintenance, a behavior consistent with asymmetric fate in which the size of the labelled NSC pool remains constant over time while neuronal progeny is produced at a constant rate.

Such fate asymmetry can be achieved either at the level of each and every NSC division or at the level of the population. Therefore, to gain insight into the fate of individual NSCs, we monitored the behavior and composition of *her4+* NSC-derived clones over the ~ 17 month chase period. We first noted that NSCs (Gs^+^ Sox2^+^) maintained an overall fixed ratio to NPs (Gs^−^ Sox2^+^) whether considering the whole population of pallial progenitors or the *her4* lineage (Fig. S8). These observations suggest that the behavior of the entire population of Sox2^+^ pallial progenitors is slave to that of the renewing NSC population. Therefore, since Sox2 nuclear localization improves the reliability of cell identity assignment, Sox2 expression was used as a surrogate NSC marker. Consistent with a population asymmetry-based model of NSC self-renewal, we found that the fraction of clones containing at least one NSC (hereafter referred to as “active clones”) decreased continuously over the first several months post-induction, and was matched by a concomitant increase in the average number of NSCs per active clone (Figs. 2A-E, S9, S10A,B). However, by 183 dpi, both measures reached a plateau that persisted until the end of the analysis. Importantly, the size of the traced NSC population (Fig. 1F,G), the proliferative activity of NSCs, as well as their fate choices at division remained approximately constant over the chase period (Fig. S10C,D, S12D), indicating that the apparent crossover in clone behavior was not associated with a change in developmental dynamics. The emergence of a plateau in clone statistics stands in contrast to that expected for a continuous process of NSC loss and replacement, where “neutral drift” would result in an ever-decreasing clone number (49), and suggests instead a model in which long-time renewal potential is invested in a subpopulation of NSCs that adopt asymmetric fate at the level of individual cell divisions. Thus, we hypothesized that *her4.1* -expressing NSCs were organized in a proliferative hierarchy, with a self-renewing subpopulation of “reservoir NSCs” (rNSCs) supporting a second population of “operational NSCs” (oNSCs) with a definite average neurogenic potential.

**Fig. 2.**
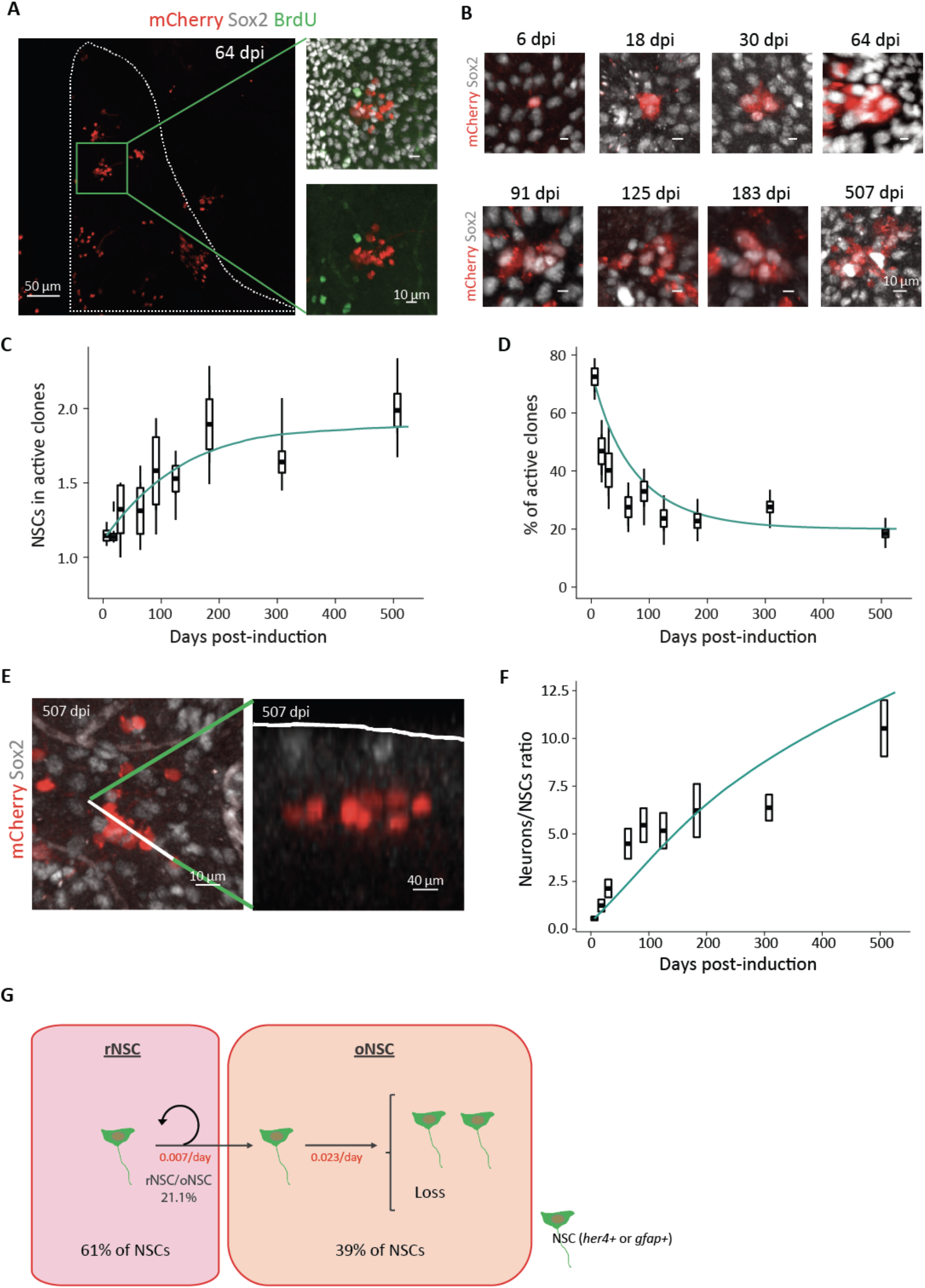
Hierarchical organization of pallial NSCs. **(A)** Dorsal view of a pallial hemisphere exemplifying some clones recovered at 64 dpi (magnification of the boxed area shows one clone). The analyzed region is outlined with dotted lines. (B) Three-dimensional reconstruction of clones at different time points illustrating the progressive increase in their NSC content until 183 dpi. **(C)** Time evolution of the number of NSCs (i.e. Sox2^+^ cells) per active clone. **(D)** Time evolution of the proportion of active clones (i.e. clones containing at least one NSCs). **(C, D)** Box and whisker plots: experimental data. The central bold bar and the upper and lower edges of the boxes represent respectively the mean and s.e.m. of the most likely clonal composition; the whiskers of the box correspond respectively to the 95% CI of the smallest and biggest clustering that are still within the 95% CI of the most likely clustering (see supplementary text). They reflect the combined uncertainty stemming from the clonal reconstruction and the finite sample size. **(E)** Left: three-dimensional (3D) reconstruction of a multi-celled fully neuronal clone at 507 dpi. Right: Optical section along the plane defined by the dotted line on the 3D reconstruction (left). Note that the clone does not contain NSCs anymore and is completely detached from the ventricular zone (dotted line). **(F)** Time evolution of the ratio of neurons to NSCs in all traced clones. **(G)** Schematic illustrating the simplified model used for MLE. The inferred values of the parameters are given in red (frequency of events per cell; rNSC ➔ rNSC+oNSC: v=0.007d^−^; oNSC ➔ oNSC+oNSC: λ=0.006d^−1^; oNSC➔loss: μ=0.017d^−1^). **(C, D, F)** Clones as defined by the algorithm. n= 6, 3, 3, 3, 5, 6, 7, 8 and 6 brains at 6, 18, 30, 64, 91, 125, 183, 307 and 507 dpi, respectively. Solid lines depict modelling predictions.

To test whether such a scheme could describe quantitatively the clonal data, we developed a minimal model. Within this framework, rNSCs divide asymmetrically at rate v, giving rise to oNSCs. To capture the variability of clonal outputs, we proposed that oNSCs follow a pattern of stochastic fate, duplicating at rate λ and becoming lost through differentiation at rate μ (Fig. 2G). (Later, we consider a more refined model that takes into account the different channels of individual oNSC fate, viz. the relative frequencies of symmetric and asymmetric divisions.) From the frequency of active clones at long times (Fig. 2D), it followed that some 20% of initially-labeled NSCs belong to the reservoir pool. A fit to the data using maximum likelihood estimation indicated that, within the endogenous NSC population, rNSCs actually constitute some 61% of NSCs and enter into cycle around once every 143 days, on average, while oNSCs (forming the remainding 39% of NSCs) select between duplication or loss once every 43 days in the ratio of 1 to 3, respectively (for details and statistical confidence intervals see Fig. 2G and S11A, supplementary text). Notably, in addition to the average NSC content and persistence of active clones (Fig. 2C,D), the model could predict the detailed clone size distributions over the entire time course (Fig. S11B,C). Although NSCs and NPs were pooled to increase statistical confidence, quantitatively similar results were obtained when the analysis was performed on the GS^+^ compartment alone.

To further test the basis of this hierarchical scheme, and dissect in detail the fate behaviors of NSCs, we took advantage of our intravital imaging method, which permits individual NSCs to be recorded in their niche, tracking >300 NSCs per hemisphere in the Dm over several weeks under fully non-invasive conditions (20). *gfap:dtomato* transgenic individuals were crossed into the transparent *casper* background (32) and imaged live over 23 days (Fig. 3A) (20). Analysis of >80 tracks of NSCs that became active showed that the fate choice of most had become revealed by change in marker expression (loss of Gfap) by around 10 days after division (Fig. S12A,B). Consequently, tracks with less than 10 days available after division were not scored (Fig. S12C). Based on this cohort, we found that some 35.8±0.7% of NSC divisions result, on average, in symmetric amplifying fate (NSC/NSC fate, as assessed by Gfap expression), 53.1±2.5% result in asymmetric fate (NSC/n, where “n” is a neuron or a NP), and 11.1 ± 1.6% result in symmetric differentiative fate (n/n) (Fig. 3B,C). Similar statistics were inferred from the analysis of two-cell clones recovered at 6 dpi, with values that were not altered with age (Fig. S12D). Notably, intravital imaging also revealed the occurrence of direct neuronal differentiation events where Gfap expression was lost without cell division (Fig. 3D and S12C,E) (18). In line with the homeostatic nature of the cell dynamics within the *her4.1* lineage (Fig 1F,G), we found that direct neuronal differentiations accounted for about 20.2 ± 1.7% of all fate choice events, thus contributing significantly to the overall balance in fates (Fig. 3E) (18)(20).

**Fig. 3.**
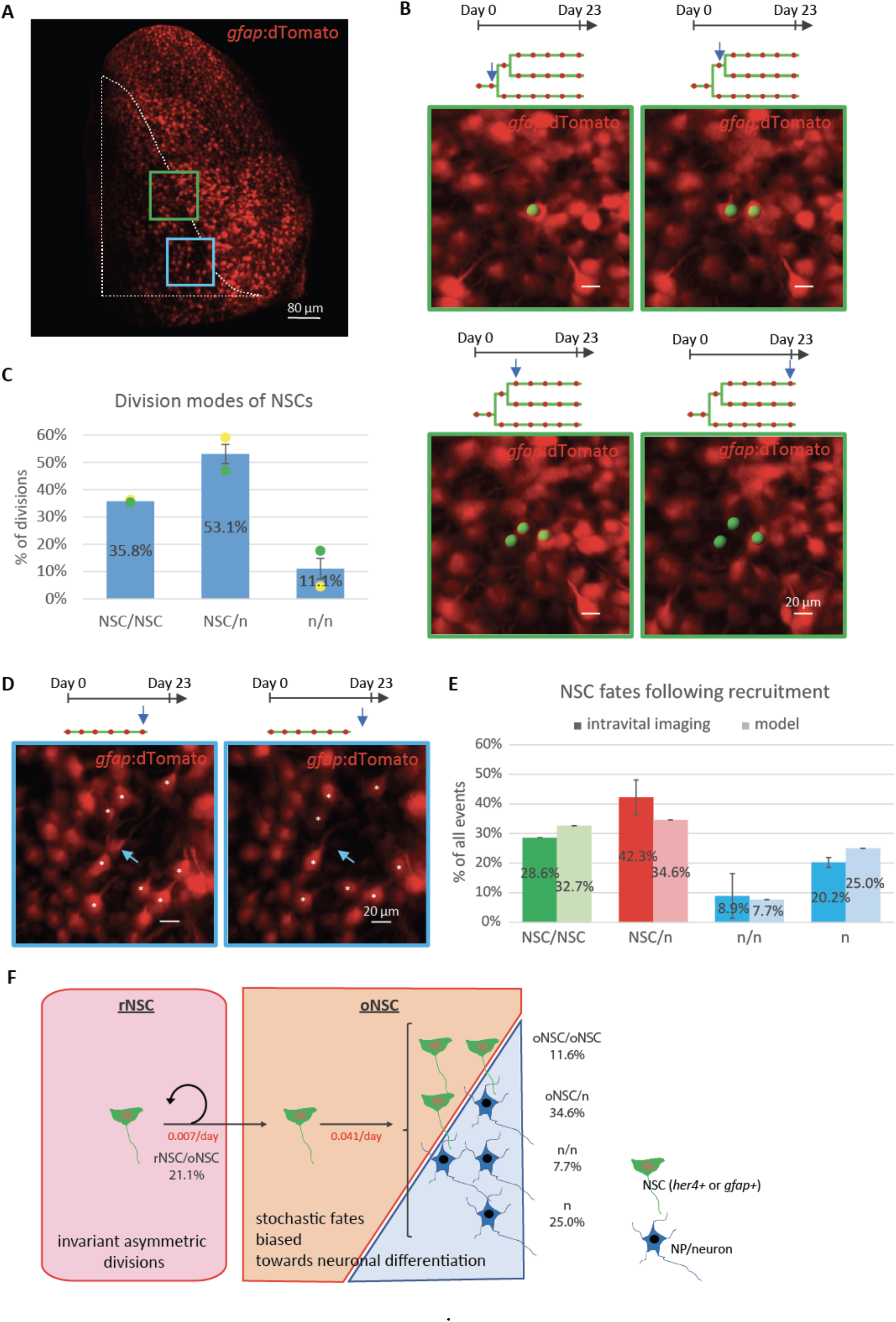
Time-lapse intra-vital imaging analysis of NSC division patterns. **(A)** Dorsal view (3D reconstruction) of a pallial hemisphere imaged from a live *casper;gfap:dTomato* fish. The region analyzed is surrounded by the dotted line. The green and blue squared areas are displayed in B and D, respectively. **(B)** Example of a NSC that underwent two symmetric amplifying divisions. Green spots are superimposed to the NSCs of interest and tracks (green) with red dots indicate imaging sessions where the NSCs were still present (blue arrows point to the moments of image acquisition). **(C)** Relative proportions of the different types of NSC divisions inferred from their tracking in live fish (color-coded). n: loss of *gfap* expression (NP/future neuron). 2 brains analyzed. **(D)** Example of a direct differentiation of a NSC (blue arrow). * mark surrounding quiescent NSCs as visual landmarks. **(E)** Relative proportions of NSC fates leading to expansion (symmetric amplifying divisions, green), maintenance (asymmetric divisions, red) and consumption (symmetric neurogenic divisions and direct neuronal differentiations, blue). Light colors are the results of our simulations. Note the overall balance in NSC fates. n=2 brains. Error bars: s.d. and credibility interval (see supplementary theory) for the *in vivo* data and the modelling, respectively. **(F)** Modelling of NSC dynamics. The inferred proportions and fate frequencies (in red, values per cell) of both reservoir and operational NSCs among all NSC activation events are indicated (oNSC➔oNSC+n: ĸ=0.018d^−1^; oNSC➔n+n: μ1=0.004d^−1^; oNSC➔n: μ2=0.013d^−1^). The corresponding curves are plotted in green above experimental data in panels C, D and F of figure 2. Note that NSC/NSC fates monitored by live imaging include rNSC/oNSC and oNSC/oNSC divisions.

To comprehensively model the *her4.1* lineage, we then introduced the different qualitative fates observed by intravital imaging (Fig. 3A-D) into the previously determined dynamical parameters of r,oNSCs (Fig. 2G), to infer the predicted rates of both asymmetrical (*κ*) and symmetric neurogenic divisions (μ_1_) of oNSCs. This allowed us to provide a complete description of NSC dynamics, including the rates of all fate transitions for oNSCs (Fig. 3F). As an important validation, we found the quantitative pattern of NSC fates predicted by our model to be in very good agreement with the measurements scored in live fish (Fig. 3E) (“NSC/NSC” fates include the divisions of rNSCs -which are asymmetric and generate rNSC/oNSC fate- and the amplifying divisions of oNSCs -oNSC/oNSC fate-, which are indistinguishable in live-imaging). Furthermore, the derived frequencies of all NSC divisions (rNSCs and oNSCs) add up to 0.035 per day, a figure commensurate with the measured division rate of pallial NSCs (Fig. S1D, S10C).

Overall, these results indicate that pallial NSCs are heterogeneous and organized in a hierarchy in which an upstream reservoir population divides asymmetrically – ensuring lineage homeostasis – and infrequently to produce operational NSCs with only limited renewal potential. The latter, despite also being mostly quiescent, activate more frequently and follow a pattern of stochastic fate biased towards neuronal differentiation, thus comprising the effective neurogenic pool.

The perfectly homeostatic behavior of the *her4.1:iCre* traced NSC population was unexpected. Since the zebrafish brain continues to grow during adulthood, expansion of the NSC pool has been long-hypothesized to accompany the ongoing growth of the pallium. Accordingly, we found that the total number of pallial NSCs expanded roughly linearly with time until 9 months of age (Fig. 4A, and S13A). Analysis of nearest neighbor distances (NNDs) revealed no significant change in NSC density over the same time frame, as well as no correlation between NSC numbers and densities, suggesting that the ventricular area progressively enlarges as the brain and the NSC pool grow (Fig. S13B,C). Variation in labelling efficiency did not blur a potential increase in the traced NSC pool as we found no correlation between the number of *her4.1:iCre-traced* NSCs and the total number of NSCs, and only a poor and non-relevant correlation between the total number of NSCs and the NSC content of active *her4.1:iCre*-derived clones (Fig. 4B and S13D). Furthermore, during the growth period, the *her4.1:iCre-derived* pool of NSCs tended to represent an ever-decreasing fraction of the total NSC population (Fig. S13E), as expected for a homeostatic pool evolving in a continuously growing population. As *her4.1^+^* and Sox2^+^ cells maintained a fixed ratio with time, it follows that the overall population of *her4.1^+^* NSCs continued to expand (Fig. 4A and S13F-H).

**Fig. 4.**
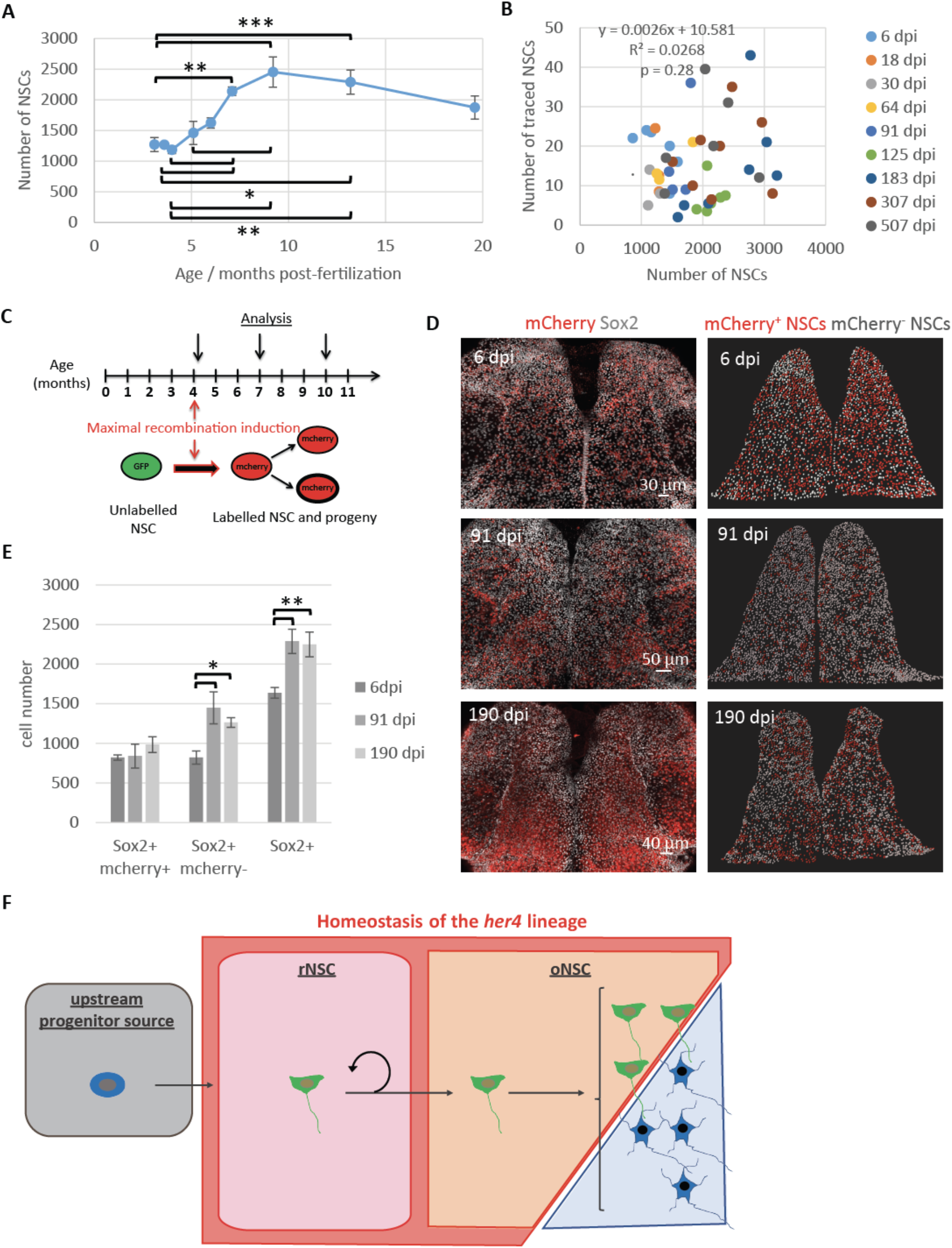
Ongoing production of her4^+^ NSCs by an upstream progenitor source. **(A)** Corresponding quantification. One-way ANOVA: F_(8,37)_=8.06, p<0.001; all pairwise comparisons: LSD test followed by Holm’s adjustment. * p < 0.05, ** p<0.01, ***p<0.001. n= 6, 3, 3, 3, 5, 6, 7, 8 and 6 brains at 3.1, 3.6, 4, 5.1, 6, 7.1, 9.2, 13.2 and 19.6 mpf (months post-fertilization), respectively. Error bars: s.e.m. **(B)** Scatter plot showing the absence of correlation between the total number of NSCs in the pallial region analyzed and the number of NSCs in the traced *her4* lineage. n= 6, 3, 3, 3, 5, 6, 7, 8 and 6 brains at 6, 18, 30, 64, 91, 125, 183, 307 and 507 dpi, respectively. **(C)** Time line of the experiment. Maximal recombination of the ubi:Switch reporter was induced by treating *her4.1*:iCre;ubi:Switch transgenic fish with 4-OHT repetitively over 5 days. **(D)** Left: Dorsal view of pallia immunostained for mCherry and Sox2 at 6, 91 and 190 dpi. Right: Segmented Sox2^+^ NSCs (spots). Red spots: NSCs within the *her4.1* lineage (Sox2+,mCherry^+^); white spots: mostly newly formed NSCs (Sox2^+^,mCherry^−^). **(E)** Quantification of Sox2^+^, mCherry^+^, Sox2^+^,mCherry^−^ and Sox2^+^ cells in the Dm territory of interest. Oneway ANOVA: Sox2^+^, mCherry^+^ cells: F_(2,13)_=0.59, p=0.5697; Sox2^+^, mCherry^−^ cells: F_(2,13)_=6.02, p=0.0141; Sox2^+^ cells: F_(2,13)_=9.06, p=0.0034. All pairwise comparisons: LSD test followed by Holm’s adjustment. * p< 0.05, ** p<0.01. Error bars: s.e.m. n= 5, 6, and 5 brains at 6, 91 and 190 dpi, respectively. **(F)** Proposed hierarchical organization of pallial neural progenitors.

Together, these results suggest that new *her4.1^+^* NSCs are generated over time from initially unlabeled cells, i.e. which do not belong to the *her4.1^+^* lineage, as targeted by the clonal induction. To add direct support to this interpretation, we devised a protocol for maximal induction with the *her4.1:iCre* driver line (Fig. 4C), initially labeling around 50% of Sox2^+^ cells (~ 70% of *her4.1^+^* cells, Fig. 4D,E and S1B). Quantification of the traced (mCherry^+^) NSC bulk population confirmed the invariance of the number of NSCs in the *her4.1^+^* lineage (Fig. 4D,E and 1F,G). In striking contrast, quantification of unlabeled (mCherry^−^) NSC population revealed their massive expansion during the chase time (Fig. 4D,E), implying that new NSCs are continuously produced within the pallial niche. Hence, our results point to the existence of an as yet undefined “NSC source”, hierarchically upstream of *her4.1^+^* NSCs, and responsible for their continuing production over time. The dynamics of the whole NSC population also predicts that the activity of the source fades after ages post 9-10 mpf (Fig. 4A).

Finally, we took advantage of the clonal data to gain insight into the production dynamics of new neurons. The total number of labelled neurons progressively increased over the full extent of the experiment which, with regard to the constant number of traced NSCs, indicates that adult-born neurons accumulate over time (Fig. 1F,G, 4E and S14A,B). In agreement with the near-absence of apoptosis during adult neurogenesis in this pallial region (19), our quantitative model also provided a very good prediction of the time increase in the ratio of neurons to NSCs in the lineage (Fig. 2F). These results also applied at the level of individual clones (Fig. S14C-E and S15A). Notably, the continuous increase of the ratio of neurons to NSCs in active clones underscores that the main driver of clonal growth is the generation of new neurons (Fig. S14D-F and S15B). Further, the lack of a major inflection in the time evolution of this ratio provided additional evidence that the plateaus observed in the clonal NSC statistics were not linked to an increase in NSC quiescence. Thus, similarly to the rodent DG (33), we found that zebrafish adult pallial neurogenesis is additive, i.e. results in a net addition of new functional neurons with time (19).

Understanding the functional behavior and hierarchical organization of the NSC lineage is a prerequisite to defining their molecular identity, regulation, evolution with age, and their response to challenge. By integrating genetic tracing, intravital imaging and global population assessments at short- and longterm over a lifetime, our results provide evidence that, in the adult zebrafish pallium, slow-cycling self-renewing NSCs (termed reservoir NSCs) reside at the apex of a proliferative hierarchy and divide asymmetrically to give rise to NSCs (termed operational NSCs) with only limited renewal potential and characterized by stochastic fate. Above this hierarchy lies an as yet uncharacterized source that fuels the continuous production of an intrinsically homeostatic population of NSCs (Fig. S4F).

By providing the first complete model of NSC maintenance in the adult vertebrate brain based on comprehensive quantitative dynamics, this study offers important conclusions: First, it demonstrates that common molecular and cellular signatures of NSCs *(her4* and *gfap* expression, and radial glia nature) mask functional heterogeneities. Second, it shows that NSC dynamics involves a hierarchy of specialized sub-populations, comprising *bona fide* self-renewing NSCs (reservoir) supporting a shorter-lived (operational) neurogenic pool. As a result, this model may unite previous phenotypical or lineage work on mammalian adult NSC pools. A partition of NSCs between a mostly dormant and a more active pool (in which NSCs shuttle between rest and activation) was recently proposed in the mouse DG based on mutant analyses (34). Likewise, *Hes5*-expressing DG NSCs as well as Troy-expressing SEZ NSCs also appear to conform to a homeostatic behavior (35), raising the possibility that the previously reported activity-dependent consumption of NSCs might rather reflect the dynamics of an operational/activated pool (8, 9, 12, 14, 36). Finally, most intriguingly, NSCs within the *Nestin* lineage of the DG were also shown to increase in number over time (17), suggesting that a source population responsible for NSC production might also exist in the mouse brain. Thus, while the model we propose primarily applies to the adult pallial neurogenic area of the zebrafish brain – potentially reconciling apparently conflicting results (18, 19) – it may provide an interesting framework to rethink and explore further NSC self-renewal and diversity in other systems, such as the mammalian brain. It also suggests that adult germinal populations that increase in cell number during adult life may combine homeostatic dynamics typical of mature tissues with a generator (“source”) akin to embryonic growth zones.

## Supporting information

All supplementary figures

## Acknowledgments

We thank members of the ZEN team for their constant input, David Morizet for advice on R usage, and Sébastien Bedu together Nicolas Chanthapathet for expert fish care. We are indebted to Dr. Shahragim Tajbakhsh for discussions and advice at initial stages of this work. Work in the L. B-C. lab was funded by the ANR (grant ANR-2012-BSV4-0004-01, and Labex Revive), Centre National de la Recherche Scientifique, Ecole des Neurosciences de Paris (ENP), Institut Pasteur and the European Research Council (AdG 322936). E. T-T was recipient of a PhD student fellowship from the Ministry of Science and Education and the Fondation pour la Recherche Médicale (FRM). B.D.S also acknowledges funding from the Royal Society E.P. Abraham Research Professorship (RP\R1\180165) and Wellcome Trust (098357/Z/12/Z).

## Materials and Methods

### Zebrafish care and strains

All procedures relating to zebrafish (*Danio rerio*) care and treatment conformed to the directive 2010/63/EU of the European Parliament and of the council of the European Union. Zebrafish were kept in 3.5-liter tanks at a maximal density of five per liter, in 28.5°C and pH 7.4 water. They were maintained on a 14 hours light / 10 hours dark cycle (light was on from 8 am to 10 pm), and fed three times a day with rotifers until fourteen days post-fertilization (14 dpf) and with standard commercial dry food (Gemma Micro from Skretting*) afterwards. All transgenic lines – *Tg(her4.1:dRFP) (37), Tg(gfap:nGFP)^mi2004^ (38), Tg(her4.1:ERT2CreERT2)* (referred to as *her4.1:iCre)* (39) and *Tg(−3.5ubb:loxP-EGFP-loxP-mCherry) (referred to as ubi:Switch)* (40) – were maintained on an AB background. The *Tg(gfap:dTomato)* (41) transgenic fish were kept on a *Casper (roy^−/−^;nacre^−/−^)* background (32). *Tg(her4.1:dRFP)/+* fish were interbred with *Tg(gfap:nGFP)/+* fish to obtain *Tg(her4.1:dRFP)/+; Tg(gfap:nGFP)/+* double transgenic fish. Similarly, crosses between *Tg(her4.1:ERT2CreERT2)/+* and *Tg(−3.5ubb:loxP-EGFP-loxP-mCherry)/+* zebrafish yielded the *her4.1:iCre; ubi:Switch* double transgenic individuals used for the lineage tracing experiments. Ages of the fish are explicitly stated in the respective experiments (see also, 4-hydroxytamoxifen treatment), except for the *Tg(her4.1:dRFP)/+*; *Tg(gfap:nGFP)/+* and *Tg(her4.1:ERT2CreERT2)/+; Tg(her4.1:dRFP)/+* double transgenic fish as well as the Casper *Tg(gfap:dTomato)* fish, which were all four-month post-fertilization (4mpf). Fish were euthanized in ice-cold water (temperature comprised between 1° and 2°C) for ten minutes, according to a special dispensation and following the guidelines of the Ministry of superior education, research and innovation.

### Genotyping

Fish were screened for the expression of the *her4.1:dRFP*, the *gfap.nGFP* and the *ubi:Switch* transgenes between 48 and 72 hpf. The presence of the *her4.1:ERT2CreERT2* transgene was detected by PCR amplification of a part of the ERT2 sequence on a tail DNA sample. DNA was extracted with the “phire animal tissue direct PCR kit” (Thermo Scientific) according to the manufacturer instructions and PCR was performed with the following primers (39):

- forward 5’-GACCCTCCATGATCAGGTCCACC-3’,
- reverse 5’-GACCGTGGCAGGGAAACCCTCTG-3’.

The thermocycling parameters were as follows:

| Cycle step | Temperature | Time | Cycles |
| --- | --- | --- | --- |
| Initial denaturation | 94°C | 2 min | 1 |
| Denaturation | 94°C | 15 s | 30 |
| Annealing | 58.5°C | 45 s |  |
| Extension | 72°C | 45 s |  |
| Final extension | 72°C | 1 min | 1 |
|  | 4°C | Hold |  |

The resulting 676 bp amplicon was resolved on a 2% agarose gel containing 0.002% SYBR safe (ThermoFisher, S33102).

### 4-hydroxytamoxifen treatment

*her4.1:iCre; ubi:Switch* double transgenic fish were immersed either for ten minutes in fish water containing 0.5 μM 4-hydroxytamoxifen (4-OHT - Sigma-Aldrich, T176) for clonal recombination of the reporter construct or 7 hours per day, for five consecutive days, in fish water containing 2 μM 4-OHT for maximal recombination. Practically, fish were placed in beakers containing 4-OHT dissolved in fish water and protected from light with foil. We used 25 ml of solution per fish for the 10 minutes induction protocol and 100 ml per fish for the full induction protocol. For the latter, fish were transferred to fresh water every night and fed in the morning before being returned to 4-OHT-containing water; water and 4-OHT solution were continuously oxygenated through a pomp-generated airflow and the 4-OHT solution was renewed every day. In both protocols, fish were rinsed at least 3 times with fresh water over 48 hours before being taken back to the fish facility. All fish used for the clonal analysis were 3 to 3.5 mpf at the time of induction. The analysis of two-celled clones (doublets of induced cells) in young and old fish was performed six days after induction in 3 mpf fish and in 14 mpf fish, respectively. The full induction experiment was carried out with 4 mpf fish.

### BrdU treatment

Proliferating progenitors of both *her4.1:dRFP* and clonally induced *her4.1:iCre; ubi:Switch* fish were labelled by a 24-hour 5-bromo-2’-deoxyuridine (BrdU) pulse, just before being sacrificed. For this purpose, fish were placed into fish water containing 1 mM BrdU and 0.333% of dimethyl sulfoxide (DMSO – Thermo Scientific, 20688). Treatments were performed in beakers, with a maximum of 10 fish per liter of solution, under continuous air flow and at a temperature of 28°C. Fish were rinsed two times five minutes in fresh water before sacrifice and dissection.

### Anesthesia (live imaging)

Anesthesia was initiated by soaking the fish for approximatively 90 seconds in water containing 0.01% MS222 (Sigma). They were then transferred into a water solution of 0.005% (v/v) MS222 and 0.005% (v/v) isoflurane to maintain the anesthesia during the whole duration of the imaging session (20). Overall, fish were anesthetized for about 30 minutes per session.

### Histology

Brains were dissected in phosphate buffered saline (PBS – Fisher Bioreagents) and directly transferred to a 4% paraformaldehyde solution in PBS for fixation. They were fixed either for 2 hours at room temperature or overnight at 4°C under permanent agitation. After two washing steps in PBS, brains were dehydrated through a series of 25%, 50% and 75% methanol in PBS and kept in 100% methanol (Sigma-Aldrich, 322415) at −20°C. Following rehydration, brains were processed for whole-mount immunohistochemistry. An antigen retrieval step was performed for subsequent BrdU and/or Pcna immunolabelling. For BrdU, brains were incubated in 2M HCl (258148 – Sigma-Aldrich) at room temperature for 30 minutes whereas for Pcna immunolabelling they were incubated with Histo-VT One (Nacalai Tesque) for an hour at 65°C. Brains were rinsed three times for at least five minutes in PBS and then blocked with 4% normal goat serum, 0.1% DMSO and 0.1% Triton X-100 (Sigma Life Science – 1002135493) in PBS (blocking buffer). They were then incubated at room temperature for 24h in primary antibodies (table 1) diluted in blocking buffer, rinsed three times over 24 hours at room temperature with 0.1% tween-20 (Sigma Life Science – P9416) in PBS (PBT) and incubated in secondary antibodies (table 2) for another 24 hours at room temperature. After at least three rinses in PBT over 24 hours, brains were transferred into PBS and their telencephali were dissected out. Every washing and incubation step was performed on a rocking platform and, from the secondary antibodies onwards, was carried out protected from light. Dissected telencephali were mounted in depression slides with Aqua Poly/Mount (Polysciences, 18606). The brain sections presented in figure S15 were processed similarly, with the following modifications. After rehydration, brains were embedded in 3% agarose, sliced in 50 μm-thick sections with a vibratome (Leica, VT1000 S) and recovered in PBS. Sections were incubated with primary and secondary antibodies overnight at 4°C and rinsed three times over the day in PBT. Finally, sections were incubated for ten minutes in 1 μg/ml 4’,6-diamidino-2-phenylindole (DAPI) in PBS, rinsed three times five minutes in PBS and mounted on slides with Aqua Poly/Mount.

| Primary antibodies |  |  |  |  |  |
| --- | --- | --- | --- | --- | --- |
| Antigen | Species | Isotype | Dilution | Source | Reference |
| Glutamine synthetase (Gs) | Mouse | IgG2a | 1:1000 | Merck | MAB302 |
| SRY (sex determining region Y)-box 2 (Sox2) | Mouse | IgG1 | 1:200 | Abcam | ab171380 |
| DsRed | Rabbit |  | 1:250 | Takara | 632496 |
| 5-Bromo-2'-deoxyuridine (BrdU) | Rat | IgG2a | 1:150 | Abcam | ab6326 |
| Proliferating cell nuclear antigen (Pcna) | Mouse | IgG2a | 1:500 | Santa Cruz Biotechnology | PC10 |
| Green fluorescent protein (GFP) | Chicken | IgY | 1:1000 | Aves Labs | GFP-1020 |
| Estrogen Receptor alpha (ERα) | Rabbit | IgG | 1:200 | Abcam | ab16660 |
| mCherry | Chicken |  | 1:1000 | EnCor Biotechnology | CPCA-mCherry |

| Secondary antibodies |  |  |  |  |  |  |
| --- | --- | --- | --- | --- | --- | --- |
| Antigen | Species | Isotype | Conjugate | Dilution | Source | Reference |
| Chicken IgY (H+L) | Goat | IgG | Alexa Fluor 488 | 1:1000 | Invitrogen | A-11039 |
| Rabbit IgG (H+L) | Goat | IgG | Alexa Fluor 546 | 1:1000 | Invitrogen | A-11010 |
| Mouse IgG2a | Goat | IgG | Alexa Fluor 633 | 1:1000 | Invitrogen | A-21136 |
| Rabbit IgG (H+L) | Goat | IgG | Alexa Fluor 405 | 1:1000 | Invitrogen | A-31556 |
| Mouse IgG1 | Goat | IgG | Alexa Fluor 647 | 1:1000 | Invitrogen | A-21240 |
| Rat IgG (H+L) | Goat | IgG | Alexa Fluor 488 | 1:1000 | Invitrogen | A-11006 |
| Mouse IgG2a | Goat | IgG | DyLight 405 | 1:1000 | BioLegend | 409209 |
| Mouse IgG2a | Goat | IgG | Alexa Fluor 488 | 1:1000 | Invitrogen | A-21131 |
| Rabbit IgG (H+L) | Goat | IgG | Alexa Fluor 488 | 1:1000 | Invitrogen | A-11008 |

### Image acquisition

Images of both whole-mounted telencephali and sections were acquired on confocal microscopes (LSM700 and LSM710, Zeiss), using either a 20 X air objective or a 40 X oil objective (for the comparison of *gfap*:nlsGFP and *her4*.: dRFP expression). With the 20X air objective, we used a z-step of 1 μm, a numerical aperture in each channel resulting in 1.5 μm-thick optical sections and a pixel dwell time of either 1.58 or 3.15 μs. Each line was averaged twice. With the 40 X oil objective, we acquired 1 μm-thick optical sections, with a step of 0.75 μm and a dwell pixel time of 1.58 μs; no averaging was performed. The power of the lasers as well as the voltage of the photomultipliers (gain) were set for every single acquisition and adjusted for signal loss throughout tissue depth. For this purpose, lasers and gain were tuned at two to three different optical sections within a z-stack and their values were linearly interpolated and extrapolated to cover the whole thickness of the stack. Images were saved in either LSM or CZI format.

The live imaging of *gfap*:dTomato-expressing NSCs was performed on a customized commercial two-photon microscope (TriM Scope II, LaVision BioTec) equipped with a ultrafast oscillator (λ = 690–1300 nm, Insight DS^+^ from Spectra-Physics Newport). dTomato was excited with photons of 1120 nm. The fluorescent signal was collected in the backward direction and separated from the excitation wavelength by a dichroic mirror (T695lpxr, Chroma) and an interference shortpass filter (ET700SP, Chroma). Photons below and above 561 nm were separated by a dichroic mirror (Di02-R561, Semrock) and directed to the “red” detection channel equipped with a GaAsP detector (H7422-40, Hamamatsu). Before collection, the “red” fluorescence was further filtered using a bandpass interference filter (FF01 – 607/70; Semrock). To image the entire volume of interest, spanning typically 800×800×250 μm^3^ (i.e. a single brain hemisphere), we recorded mosaics consisting of 4 z-stacks arranged in a square pattern with an overlap of 10%. For each z-stack the lateral field of view was 405×405 μm^2^, the depth of imaging varied from 250 to 290 μm (starting about 250 μm below the skin), the voxel size was 0.8×0.8×2 μm^3^ and the pixel dwell time was 4.9 μs.

### Image processing and cell countings

Images were exported to the Imaris solfware (versions 8 and 9, Bitplane) and converted to the Imaris file format.

#### Lineage tracing

To circumscribe the automatic detection of cells to the dorso-medial part (Dm) of the pallium (with the exception of its most posterior part), we manually drew surfaces from the most dorsal part to the most ventral part of the pallium stack, outlining the sulcus ypsiloniformis on the lateral side of Dm. The posterior limit of these surfaces was defined by the plane perpendicular to the anterior-posterior axis of the telencephalon and tangent to the posterior tips of the sulcus ypsiloniformis. The compilation of these surfaces delineated a volume, representing our region of interest (ROI), that we used to create a new specific set of channels (for every original channel, we set the voxels outside the ROI to 0). A few neurons within the parenchyma also express Sox2 and might be unduly assigned a progenitor identity. To circumvent this issue, we created a new volume corresponding to the parenchyma of our ROI from which we deleted the Sox2 channel (by setting every Sox2 voxels to 0). Overall, this strategy allowed us to use the spot function of Imaris to automatically detect both mCherry- and Sox2-expressing cells. More generally, this function allows the manual or automatic registration of objects (such as cells) as “spots”, whose coordinates can be easily recovered.

In the clonal analysis experiment, we first ran an automatic detection (“segmentation”) of mCherry-expressing cells. Then, to distinguish between Sox2^+^ and Sox2^−^ cells among the mCherry^+^ population, we used a filter based on the median intensity of the Sox2 signal contained in 3 μm-diameter spheres centered on mCherry^+^ cells (spots). A similar automated approach was not feasible for the detection of Gs-expressing cells owing the particular cytoplasmic distribution of this enzyme. As a consequence, Gs+/Sox2^+^ NSCs were manually distinguished from Gs^−^/Sox2^+^ NPs both on 3D reconstruction of z-stacks and on digital cross-sections, and each cell was registered with a spot corresponding to their respective progenitor category. Proliferating NSCs (aNSCs) and NPs (aNPs) were subsequently selected by applying a median intensity filter on the BrdU signal included in a 3 μm-diameter sphere around their corresponding spots. Given the complete absence of parenchymal astrocytes in the zebrafish telencephalon, the production of oligodendrocytes through an independent lineage (42) and the almost complete labelling of parenchymal cells by the neuronal marker HuC/D (Fig. S16), all the other cells of the lineage were deemed neurons. Finally, the whole population of Sox2-expressing progenitors was automatically segmented and their nearest neighbor distances (NNDs) were calculated with the “spots to spots closest distance” function of Imaris. Both the positions and the NNDs of Sox2^+^ progenitors as well as the positions and identities of the traced cells were exported as excel files for further processing.

The number of Sox2^+^ cells within and outside the lineage of maximally recombined *her4.1* -expressing NSCs was determined first by automatically segmenting the whole population of Sox2^+^ cells and then by applying a filter based on the median intensity of the mCherry signal within 3 μm-diameter spheres centered on Sox2^+^ spots.

#### Analysis of NSC/progenitor markers

The analysis of the distribution of the different types of pallial progenitors was carried out within a 200 μm-side cubic region included into our ROI. As Sox2 expression characterizes every progenitor contacting the pallial ventricular zone, we started by automatically segmenting Sox2-expressing cells and we then sequentially applied distinct median intensity filters to the signals of the different markers until sorting out all the different progenitor categories.

In all experiments, the automatic cell segmentation was systematically verified on 3D reconstruction of the pallial z-stacks and corrections were manually made whenever necessary. Furthermore, colocalizations of markers were also confirmed or edited for every single cell on digital cross-sections.

The telencephalic transverse sections shown in figure S16 are displayed as single confocal planes.

#### Live imaging of NSC fates

Images recovered from live fish were analyzed as previously described (20). Briefly, the z-stacks acquired on successive imaging sessions were first converted into a single file using Imaris (Bitplane) and Fiji. Time-points were then manually aligned and intensities adjusted to correct for potential fluctuations between imaging sessions. Between 300 and 400 cells were tracked over eight time points (23 days) in the pallial Dm region. While the approach implemented here does not allow to distinguish between loss of *gfap* expression and the death of *gfap*-expressing cells, we emphasize that apoptotic cell death was previously shown to be very low during neurogenesis in the adult zebrafish pallium (19).

### Statistical analysis

To ensure the consistency of the cell quantifications, the same experimenter carried out all the cell counts for a given experiment (lineage tracings, time lapse live imaging, analysis of NSC markers…). In addition, when comparisons were done between experiments, we also made sure that the same experimenter performed all the measurements in the experiments being compared. Investigators were not blind to the time of chase nor were they to the age of fish. No computational randomization methods were used, but special attention was paid to maximize the random distribution of fish across conditions. Balanced ratios of females and males were included in the different experimental groups as much as possible. In the clonal analysis experiments, all the fish batches (except the ones used for the 6 dpi time point) arose from crosses between the same pairs; whenever feasible, fish of induced batches were collected over several time points. In the full induction lineage tracing experiment, we used a single batch of fish that were all induced together.

Data are presented as mean ± standard error of the mean (s.e.m.), as mean ± standard deviation (s.d.) or as mean ± 95% confidence interval (95% CI). Means represent averages per hemisphere per animal (the statistics of both hemispheres were averaged in such a way that we were able to include in the analysis the brains with only one hemisphere left). Statistical analyses were carried out using InVivoStat (43) and plots were created using either Microsoft Excel or R. The normality of the residuals of the responses was assessed using normality probability plots and the homogeneity of the variance was inspected on a predicted versus residual plot (44). When the responses deviated noticeably from either criterion, they were first log10-transformed. In addition, all proportion responses were transformed using the arcsine function (44). Data displaying an approximately Gaussian distribution of residuals and homoscedastic responses with or without transformation were analysed using parametric tests. When factors (e.g. time of chase, cell types) were analysed at more than two levels (6 dpi, 18 dpi, 30 dpi.; *her4.1*^+^ NSCs, *Gfap*^+^ NSCs, NPs.), overall effects were determined either by analysis of variance (ANOVA) or with a repeated measures mixed model approach, if animals were measured repeatedly. In this case, the within-subject correlations were modelled by a compound symmetric covariance structure. No gateway ANOVA approach was used and pairwise comparisons were carried out independently of the results of the ANOVA with least significant difference tests (LSD). P-values were adjusted for multiple comparisons according to the Holm’s procedure. Single comparisons were analyzed with independent (unpaired) Student’s t-tests when the variances of the factor levels were similar, with Welch’s t-tests when the corresponding variances were unequal and with a paired t-test when the two levels of the investigated factor were measured in the same animals. Responses that continued to harbor a significant deviation of their residuals from a normal distribution and/or heterogeneous variances after transformation were analyzed using non-parametric tests. In that case, overall effects were assessed with a Kruskal-Wallis test and all pairwise comparisons with Behrens Fisher tests (44, 45). Categorical (binary) responses (interhemispheric fusions) were analyzed using a Fisher’s exact test.

All the statistical tests performed were two-tailed and their significance level was set at 5% (α=0.05).

## Supplementary Text

### Brain region analyzed

The adult zebrafish telencephalon harbors a large neurogenic domain including the territories thought to be homologous to the two NSC-hosting niches of the mammalian brain (the subependymal zone of the lateral ventricle and the subgranular zone of the dendate gyrus). We decided to focus our study on the dorsal part of the pallial Dm region (Fig. S1A and S4). This region harbors numerous dorsally exposed NSCs, easily accessible for whole-mount immunohistochemistry and subsequent confocal imaging, as well as for intra-vital imaging. Furthermore, in Dm, the minimal migration of their neuronal progeny allows clones to remain compact and close to the surface which, beyond alleviating potential clonal ambiguities, also permits to capture the whole set of clones in a single acquisition without the need to resort to sectioning or tissue clearing. Finally, Dm is currently the best characterized pallial germinative area in zebrafish at the molecular level and has the advantage of harboring NSCs with a greater activation frequency (20), thus enhancing our chances to capture the clones’ dynamics in this overall relatively slow proliferating system (18, 26, 46). Relative to mammalian neuroanatomical subdivisions, Dm encompasses the neocortical area, which is neurogenic in adult zebrafish (22).

### Clonal induction

To determine the conditions of 4-hydroxytamoxifen (4-OHT) needed to generate unambiguous clones, we induced 3 mpf adult zebrafish with decreasing concentrations and exposure times to 4-OHT until reaching labelled cell densities compatible with the long-term expansion of clones, a property referred to as clonal density (Fig. 1B-D and S5). However, because clones may lose their last NSCs over time, a suitable clonal density is necessarily a compromise between starting with a sufficient number of induced cells to permit statistical inferences while still maintaining an acceptable risk of fusion between adjacent clones during their expansion. Accordingly, we chose 4-OHT parameters resulting in an average number of 20.7 ± 2.45 (mean ± s.e.m.) singly labelled cells (or cell clusters) per hemisphere at 6 dpi (Fig. S5C-E). By that time, although already 46% of the total number of clones had activated, only 1.6% of the Sox2^+^ progenitor population was labelled with mCherry, thus arguing for the clonal character of our data (Fig. S6 and S13E). The slow accumulation of mCherry protein precluded the reliable counting of labelled cells at earlier time points. We also noted a markedly higher induction rate of *her4.1* -expressing cells in the most posterior part the pallium, which was associated with an evident dissociation of the clones at later time points. For these reasons, this pallial area was excluded from the analysis based on anatomical landmarks (Fig. S4). Analysis of nearest neighbour distances (NNDs) between the centres of Sox2^+^ clones indicated that their majority lies at a distance greater than 44 μm from each other, i.e. about eight NSC diameters (Fig. S5A, B and S13B). Besides the sparse labelling of cells following induction, the clonality of our data set is also underscored by the invariable average number of clones across all the time points analysed (with the exception of the last at 507 days – see below) (Fig. S5C-E). This result indicates that neither clonal fusion nor fragmentation significantly affect our data. Finally, to guide the unambiguous assignment of labelled cells to putative clones, we developed a clustering algorithm based on the likelihood of putative clones to have arisen from a single induction event (Fig. S7 and supplementary text). Together, these preliminary controls warranted the clonal character of our data set and allowed its quantitative analysis.

Ultimately, we examined how the progenitor population targeted by the *her4.1*: iCre driver line at 6 dpi compared with the overall population of progenitors. In line with the relatively substantial number of clones that had already activated by this time (Fig. S6), we observed that a significant number of NPs had already formed in the lineage (S17A). Interestingly, the distribution of the main categories of progenitors in the lineage was overall similar to that found in the general population (Fig. S17A, B). This observation suggests that the upstream progenitor pool (“source”) that drives the growth of the NSC population represents only a small fraction of the non-astroglial pallial Sox2^+^ progenitors.

### Clonality of the last time point

The last time point of the clonal analysis (507 dpi) displayed a significant increase of its average number of clones relative to previous time points, thus raising concerns about its clonality (Fig. S6C). Indeed, this increase might reveal an under-clustering of the cells caused by a possible fragmentation of the clones. In fact, both the division of unlabeled progenitors and the generation of new unmarked neurons in the close vicinity of the traced clones can contribute to the scattering of their comprising cells. However, we argue that the main conclusions that we drew about the clonal dynamics at 307 dpi – i.e. the appearance of a plateau both in the number of NSCs per active clone and in the proportion of active clones – still hold at 507 dpi. Notably, as 507 dpi clones are mainly comprised of neurons, their fragmentation would be expected to lead to a decrease in the proportion of active clones, which was not the case. Furthermore, while the expanded neuronal content of the clones at 507 dpi might have blurred the unambiguous assignment of their cells, their NSC were scored with high confidence owing to the lower fraction of active clones at that time. Hence, based both on our high confidence in the maintenance of the plateau in the number of NSCs per active clones and on the stability of the proportion of active clones – which contradict a fragmentation of the clones-, we concluded that the clone dynamics observed at 507 dpi were reliable. They were thus included in our analysis.

### Aging

Aging of the zebrafish telencephalon has been associated with an increase in NSC quiescence as well as with sporadic discrete interhemispheric fusions (29). In addition, NSC density was suggested to remain approximatively constant with advancing age (29). In apparent contradiction with these reports, we did not find any significant increase in NSC quiescence (either within or outside the lineage) nor were we able to evidence any augmented proportion of pallia displaying interhemispheric fusions with advancing age (Fig. S10C, D and S18A-D). Conversely, we found that Sox2-expressing progenitors became sparser in old fish and that 44.4% of 20-month-old dorsal pallia displayed major signs of NSC depletion (Fig. S13B and S18E-F). Surprisingly, these age-associated NSC losses did not seem to be accounted for by a change in their division pattern (Fig. S12D).

Finally, we noticed that the growth of the NSC population ceased around nine months of age (Fig. 4A and S13H). In the absence of any noticeable change in pallial progenitor proliferation with age (Fig. S10D), this observation suggests that the upstream source of progenitors responsible for the expansion of the NSC population becomes progressively consumed over time. Thus, while the determination of the division mode of these progenitors will ultimately require the characterization of specific markers, we speculate that their fate may be biased towards NSC production.

## Supplemental Theory

In this supplemental theory we give further details on the calculations supporting the main conclusions drawn in the main text.

### Assignment of clonal origin to cells

The interpretation of lineage tracing data in the zebrafish pallium is complicated by the fact that stochastic forces originating from cell divisions in the surrounding tissue can lead to clone fragmentation and dispersion, which ultimately leads generic scaling distributions of clone sizes and an erasure of biological information (47). As neither the number of induced cells nor the rate of fragmentation are known a priori this renders the assignment of clonal progeny potentially ambiguous. By implementing a mathematical framework to analyze the spatial statistics of dispersed cells it is possible to recover the clonal origin of marked cells with known uncertainty and to unveil information on cell fate behavior of the traced population. We emphasize that this framework provides a general platform to assign the clonal provenance of marked cells in parallel contexts.

If induction events are statistically independent, the number of induced cells, *n*, in a given pallial hemisphere is distributed according to a Poisson distribution,

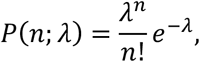

where *λ* denotes the induction frequency. Then, the distance *d* to the nearest neighbor of an induced cell in two spatial dimensions is distributed according to

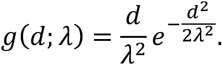

As cells divide and clones expand and disperse, the positions of labelled cells cease to be statistically independent such that the distances between nearest neighbors of labelled cells are not distributed according to *g*(*d; λ*) for any value of *λ*. However, if stochastic forces from the surrounding tissue are isotropic, the centers of marked clones defined by the average position of cells, 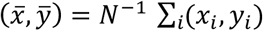, remain statistically independent and nearest neighbor distances are distributed according to *g*(*d;λ*). Therefore, for a given putative clonal assignment of cells the agreement of the distribution of nearest neighbors of clone centers with *g*(*d; λ*) can serve as a test for the correctness of this assignment, such that the true clonal assignment has the highest accordance with the assumption of statistical independence of clone centers.

To formalize this basic idea, we begin by considering the probability that a given partition of a set of cells into clones, Π, is clonal. Using Bayes theorem, we can write this probability as the product of the probability of observing the experimental data (the coordinates of labelled cells) given the clonal assignments encoded in a partition, Π, times our prior believe in a clonal partition,

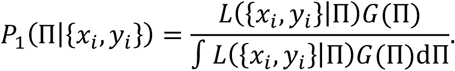

The denominator ensures normalization. The likelihood function, *L*({*x_i_,y_i_*)|Π), contains the data-dependent part of the denominator, while the prior, *G*(Π), encodes our a priori belief that a given partition is clonal. If induction events and cell fates in distinct clones are each statistically independent, the likelihood function takes the form

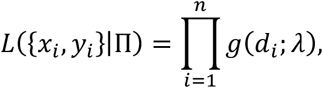

where the product goes over all putative clones in the partition Π and *d_i_* are the nearest neighbor distances of their respective centres of mass. The parameter *λ* is determined self-consistently from the number *n* of putative clones in the partition by 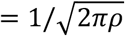, with *ρ* being the number of putative clones in the partition Π divided by the area of the analyzed tissue sample. Intuitively, the likelihood is therefore a measure for the accordance of the clonal partition Π with the hypothesis of independent labelling.

The assignment of the clonal provenance of labelled cells can be improved by considering further knowledge we might have on cell proliferation or cell migration. Specifically, in the pallium, there are biological and physical limits on cell migration, and therefore on the degree of clone dispersion. Cells are subject to stochastic forces exerted by the surrounding tissue leading to their diffusive displacement (48). If *D* is the corresponding effective diffusion constant, cell positions at a time *t* after labelling are approximately normally distributed,

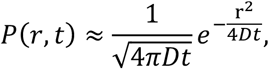

where *r* is the distance from the initial position of the induced cell. With this, we consider a tissue sample containing *n* putative clones. Then, the maximum distance to the clone centre across all clones in the tissue is obtained by calculating the n-th order statistics, i.e.

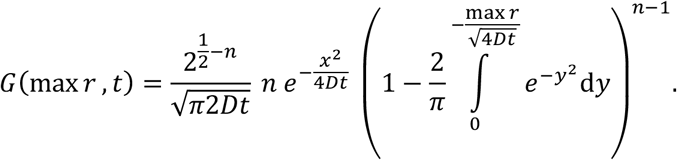

For the tail of the distribution, max 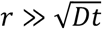, this expression is well approximated by

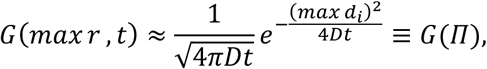

which we used as a prior in *P*_1_(Π|{*x_i_,y_i_*}). We calibrated the effective diffusion constant, *D*, by selecting a sample where clonal assignments of labelled cells could be unambiguously made by visual inspection. We then determined *D* such that the number of clones counted by visual inspection was equal to the algorithmically calculated number of clones (see below) and obtained *D* = 1.8 *μm*^2^/*d*. This means that, on average, within 100 days a labelled cell covers an area of 180 *μm*^2^ as a result of stochastic forces exerted by the surrounding tissue.

As the posterior probability trivially depends on the number of terms comprising the log likelihood, and therefore on the number of assigned clones in the partition, Π, we compare *P*_1_ to the posterior probability *P*_0_ in a scenario, where the hypothesis of statistical independence of clone centers is true. We therefore seek the clonal partition that maximize their ratio,

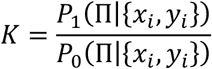

In this case, we can calculate the posterior analytically with the likelihood given by

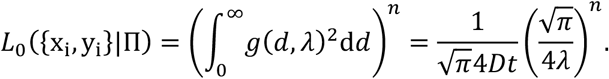

As the number of possible partitions of labelled cells into clones is too large to be computationally accessible, we employ a hierarchical or coarse-graining approach. Starting from a partition, where the number of clones is equal to the number of labelled cells, at each iteration we merge the two clones whose centers of masses have the smallest distance. While this approach bears the risk that the true clonal partition might not be tested, it allows us to obtain an approximation of this partition within only *n* iterations. The partition with the maximum value of *K* is taken for downstream analysis. To estimate the uncertainty associated with the assignment of clonality, we define all partitions whose value of *K* exceeds a threshold,

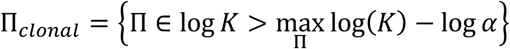

as credibly clonal.

We set *α* to 0.05 such that partitions having a value of *K* exceeding 5% of the maximum value are considered credible. We found that for all analyzed samples the mode of the posterior distribution and the corresponding credibility bounds were well defined. For each pallial hemisphere analyzed, subsequent clonal analysis was performed for the partition corresponding to the maximum value of *K* and for those corresponding to the lower and upper boundaries of the credibility interval.

### Maximum likelihood estimation

To unveil the dynamical rules that underlie NSC fate regulation in the pallium, we employ the idea of maximum likelihood estimation. Specifically, we seek to identify the model, defined by a set of parameters Θ = (Θ_1_, Θ_2_,…), that has the highest probability, *P*(Θ|*E*), given the experimental data, *E*. According to Bayes’ theorem, this probability is proportional to the probability *P*(*E*|Θ) of obtaining the experimental evidence if a certain model Θ is true while *P* (Θ) gives prior belief in the model,

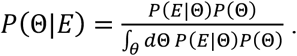

The denominator ensures normalization. The probability *P*(*E*|Θ) is called likelihood, *L_E_*(Θ) = *P*(*E*|Θ). Without prior knowledge, i.e. if the prior *P*(Θ) follows a uniform distribution, the principle of maximum likelihood states that the model with the highest probability is the one that maximizes the likelihood function, *L_E_*(Θ).

In our clonal labelling assay, the experimental evidence is given by the frequency *f_n_s_,n_n_,t_* of observations of clones with *n_s_* NSCs (“s” stands for stem cells) and *n_n_* neurons at time *t* post induction. Since the clonal observations are statistically independent, we can rewrite the likelihood function as

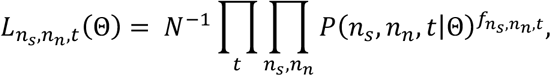

with

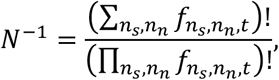

where the prefactor *N*^−1^ is determined by normalization and the products and the sums go over all time points and all clonal compositions.

In order to obtain the likelihood function, we need to calculate the probability *P*(*n_s_, n_n_, t*|Θ) to find a clonal composition with *n_s_* NSCs and *n_n_* neurons at time *t*. The time evolution of this probability is given by a master equation of the form,

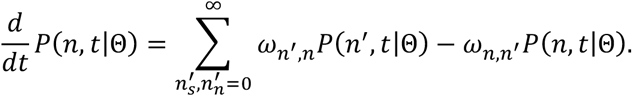

The transition rates, *ω_n,n′_*, between states with different cellular composition carry the details of the model. For simple cases the master equation can be solved analytically, however, generally, numerical methods are necessary to find approximative solutions. Using Gillespie’s algorithm, we generated stochastic trajectories whose accumulation provided histograms and consequently the probability distribution *P*(*n_s_,n_n_,t*|Θ). As extinct clones (*n* = 0) are experimentally undetectable we define the clone size distribution of active clones,

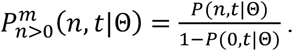

Since maximizing the likelihood function is equivalent to maximizing its logarithm, we consider the logarithm of the likelihood function,

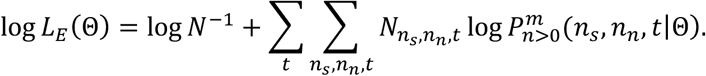

As the normalization factor *N*^−1^ is independent of the model parameters, we can neglect this prefactor in the subsequent analysis. Therefore, by substituting a particular set of experimental data in the probability distribution *P*(*n,t*|Θ), we obtain the value of the likelihood function for a given model.

### Uncertainty and credibility intervals

The finite size of the experimental and numerical samples and the variability between fish generate uncertainty in the estimation of the model parameters. To represent this uncertainty, we present the estimated parameters with margins. We define all parameters whose likelihood is above a given threshold *α* as plausible, resulting in a credibility interval,

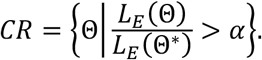

Here, we chose *α* = 0.05, meaning that parameter values, whose likelihood is more than *5*% of the value of the maximum likelihood, are considered credible. Following this, we present the value of each parameter Θ_*i*_ with its credibility interval, 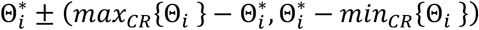.

### Inference of the best fitting model

#### NSC compartment

To begin, we focused our analysis on NSCs (Sox2^+^ cells) alone. Motivated by the observed plateau in the fraction of persisting NSC-containing clones, we hypothesized that the NSC population is heterogeneous and contains a subpopulation of long-lived reservoir NSCs (hereafter denoted as A* cells), giving rise to a downstream population of operational NSCs (referred to as A cells), whose fates are overall biased towards differentiation. In order to give rise to a plateau in the fraction of surviving clones, labelled A* need to be maintained over the time of the experiment in such a way that loss via a potential neutral drift is negligible. The saturation in the average sizes of active clones (Fig. 3C) suggests that the A* population is not downstream of A (in which case active clones would accumulate NSCs over time) but feeds into the A population. The simplest way the A* population can be maintained over long times while at the same time giving rise to cells of type A is via asymmetric cell divisions. We denote the rate with which *A*\* divides asymmetrically to give rise to both another *A*\* and an *A* cell by *v*. We note that while an asymmetrically dividing populations of reservoir NSCs is the simplest process in agreement with the clonal data, we cannot rule out a more complex cell fate behavior in this compartment, such as the existence of a closed niche or further heterogeneity.

The size of the *A* compartment can only increase by symmetric proliferating divisions or decrease due to symmetric differentiating divisions or direct differentiations. We denote the rates for both processes with *λ* and *μ*, respectively. If these operational NSCs reverted back to the reservoir NSC compartment to maintain homeostasis such a process would necessarily need to be balanced by a loss of reservoir NSCs, for example by direct differentiation to operational NSCs. Such a process would, however, lead to clonal loss due to chance fluctuations and is therefore incompatible with the clonal data. Reversion of operational NSCs therefore must be rare or nonexistent. In summary, the dynamical rules governing cell fates of the traced NSC population can be written in chemical notation as:

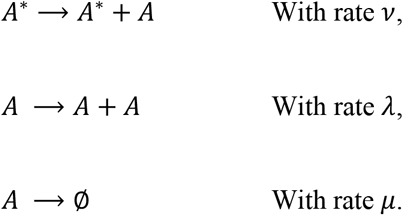

Performing maximum likelihood estimation, we obtain the best fit for parameters corresponding to a log likelihood of log *L_E_*(Θ*) = −791.55, with rates

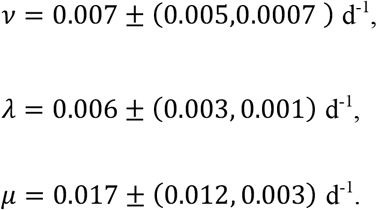

In summary, we find the best fitting parameters defining a model for a heterogeneous population of NSCs consisting of two subpopulations where, on average, *A*\*cells give rise to *A* cells every 143 ± (15.87,59.52) days, A cells duplicate every 166.67 ± (33.30,55.56) days and are lost through neuronal differentiation every 58.82 ± (12.6, 24.34) days. In the field of stochastic processes, this model is often referred to as a birth-death process with immigration (30). If *λ < μ*, as is the case here, this model ultimately gives rise to a steady state in the clone size distribution, which, in the long-term, is representative of the tissue. The best fitting parameters suggest that the traced NSC population is overall homeostatic, an observation that we were able to independently confirm by our experiments (Fig. 1E, F and 4D-F). Thus, in the steady state the ratio of A* to A cells is = (*μ* – *λ*)/*v* = 1.57 ± (0.095,0.3), such that the traced NSC population is comprised for 61% of A* cells (reservoir NSCs) and for 39% of A cells (operational NSCs).

To challenge the hypothesis of heterogeneity in the NSC population, we asked whether alternative models involving a homogeneous (equipotent) NSC population could equally explain the clonal data. First, such a model would have to predict the stochastic dynamics of active clones. The quality of such a prediction is measured by the value of the maximum of the log likelihood. In order to quantify the relative capacity of different models to describe the clonal data we estimated the relative information lost in describing the experimental data in each case, a quantity known as Akaike information criterion (AIC). Secondly, in addition to the statistics of active clones, these models need to predict the time evolution of lost clones, in particular the plateauing of the fraction of surviving clones.

In the simplest case, if the NSC population is equipotent the dynamics is determined by a simple birth-death model similar to the one proposed in other systems (49). In chemical notation,

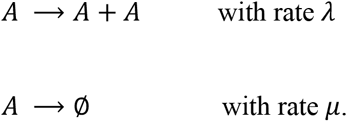

The maximum of the log likelihood is −793.2. Calculating the Akaike Information Criterion (AIC) for each of the two models (indices 1 and 2 respectively refer to the heterogeneous and the homogeneous NSC populations), we obtain *AIC*_1_ = 1589 and *AIC*_2_ = 1590. This means that, after considering the different number of parameters estimated in each case, the equipotent model is by a factor of 0.6 less likely to minimize information loss in describing the clonal data, in favor of NSC heterogeneity. While the maximum values of the log likelihoods are relatively similar, such a model containing an equipotent NSC population necessarily leads to a rapid loss of labelled clones and therefore is incompatible with the observation of a plateauing fraction of active clones over long times.

A different scenario that can result in the observed non-zero plateau with a homogeneous NSC population is a closed-niche model based on the regulation of NSC fates by their number in the niche, as suggested by Basak et al. (14) for the adult mouse sub-ependymal zone. In this model, the probability that a NSC follows either proliferation or differentiation, is coupled to the number of NSCs present in the niche,

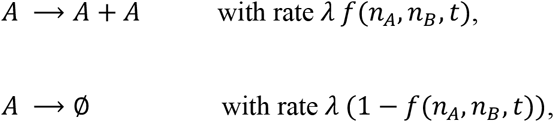

where *f*(*n_A_,n_B_, t*) = exp[-(*n_A_* + *n_B_* – *n*_0_)/*n*] is the probability of symmetric proliferation and *n_A_* and *n_B_* are, respectively, the number of labeled and unlabeled cells in the niche at time *t. n*_0_, *n* and *λ* are parameters of the model. Our simulations demonstrate that although this model can result in a plateau for the average number of NSCs in active clones, it can poorly resolve the value of the plateau in the fraction of active clones and the time scale over which it is reached. The value of the maximum log likelihood for this model is log *L_E_*(Θ*) = –817.88, meaning that the niche-based model is 1.65 × 110^−24^ fold less likely than the one comprising a heterogenous population. It is important to note that this does not rule out the existence of a closed niche governing a heterogeneous NSC population as discussed above.

To account for a potentially biased cell labelling upon induction, we initialized our simulations in such a way that clone sizes and cell type compositions resembled the clonal data from brains taken shortly after labelling (at 6dpi time). We then made predictions for all subsequent time points.

#### Neuron compartment

Having determined the dynamical rules and parameters governing cell fate behavior of the traced NSC population, we then asked whether such a model could predict the neuronal output of the labelled NSC population over time. To this end, we considered all the possible fates by which NSCs could give rise to neurons,

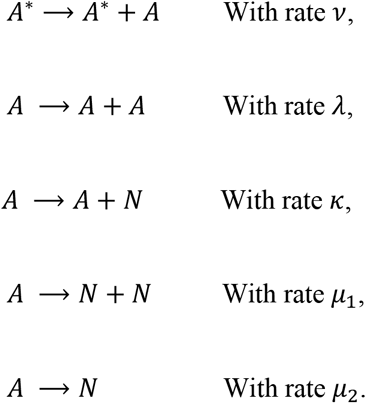

and used the maximum likelihood parameters obtained for the NSC model (*v* = 0.007 d^−1^, *λ* = 0.006 d^−1^, and *μ*_9_ + *μ*_2_ = 0.017 d^−1^). We applied maximum likelihood estimation to predict the neuronal output of operational NSCs. Using the best fitting parameters, *κ* = 0.018 ± (0.0007,0.0009) and *μ*_1_ = 0.004 ± (0.0001,0.0003), and the steady state value for the ratio of *A*\* to *A* cells (*r* = 1.57), allowed us to quantify the fractions of different fate outcomes. Specifically, the probabilities of different fate outcomes are related to the rates of symmetrical, asymmetrical, and direct differentiating events,

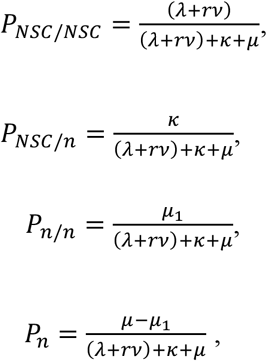

where 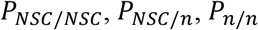, and *P_n_* are, respectively, probabilities of *NSC/NSC, NSC/n, n/n*, and *n* fate outcomes. Our analysis shows that *NSC/NSC* divisions constitute *P_NSC/NSC_* = 33± (0.05,0.1)% while *NSC/n, n/n* and *n* constitute, respectively, *P_NSC/n_* = 35 ± (0.01,0.1)%, *P_n/n_* = 7 ± (0.03,0.04)% and *P_n_* = 25 ± (0.05,0.01)% of all possible fates. As demonstrated in Fig. 3E, these percentages are in good agreement with the values obtained via our live imaging experiments.

