## Supplementary material for "Lineage hierarchies and stochasticity ensure the long-term maintenance of adult neural stem cells": All supplementary figures

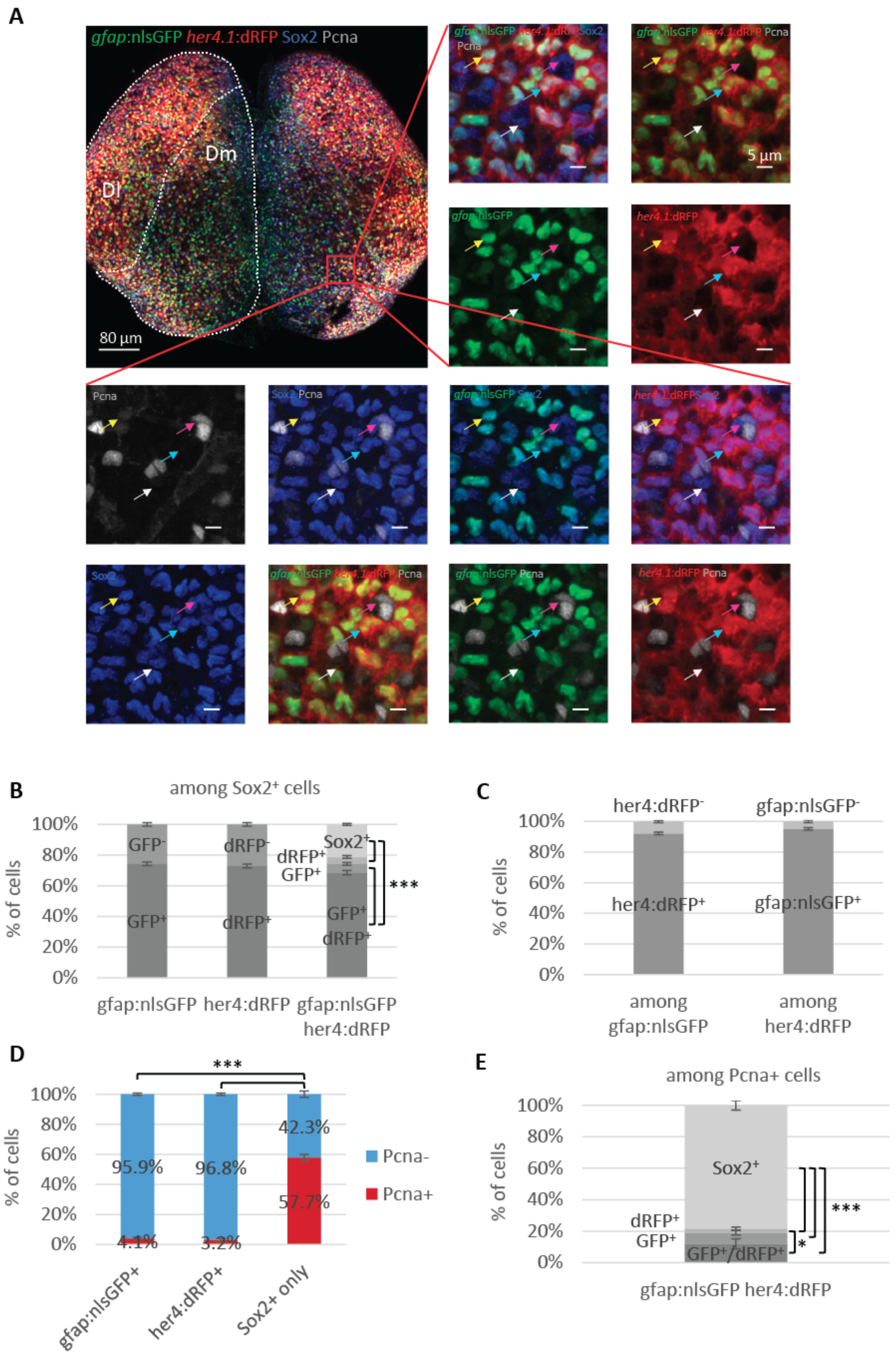


Figure S1

**Fig. S1. *her4.1*:dRFP and *gfap*:nlsGFP expression characterize the same subpopulation of Sox2-expressing pallial progenitors.** (**A**) 3-dimentional (3D) reconstruction (dorsal view) of the pallium of a *her4.1:dRFP/gfap:nlsGFP* double transgenic adult immunostained for GFP, dRFP, Sox2 and the proliferation marker Pcna. The lateral and the medial pallial domains (Dl, Dm) are highlighted with dotted lines. Close-ups of the boxed area with different combinations of channels illustrating the four Sox2$^+$ progenitor subtypes that make up the pallial germinal zone: quiescent NSCs (qNSCs – blue arrow), activated NSCs (aNSCs – yellow arrow), qNPs (white arrow) and aNPs (pink arrow). (**B**) Distribution of *gfap*$^+$ and *her4.1*$^+$ NSCs, alone and in combination, among Sox2$^+$ progenitors. (**C**) Respective distributions of *her4.1*$^+$ and *gfap*$^+$ cells among the *gfap*$^+$ and the *her4.1*$^+$ populations. Paired t-test: p=0.34 (**D**) Relative proportions of quiescent (Pcna$^-$) and proliferating (Pcna$^+$) *gfap*$^+$ NSCs, *her4.1*$^+$ NSCs, and NPs (Sox2$^+$ only). (**E**) Distribution of *gfap*$^+$ and *her4.1*$^+$ NSCs together with Sox2$^+$ NPs among proliferating (Pcna$^+$) progenitors. (**B-E**) n=7 brains analyzed. Error bar: s.e.m. (**B, D-E)** The data were analyzed using a repeated measures mixed model. Overall tests: p<0.001; pairwise comparisons: * p<0.05, *** p<0.001 after Holm's adjustment.

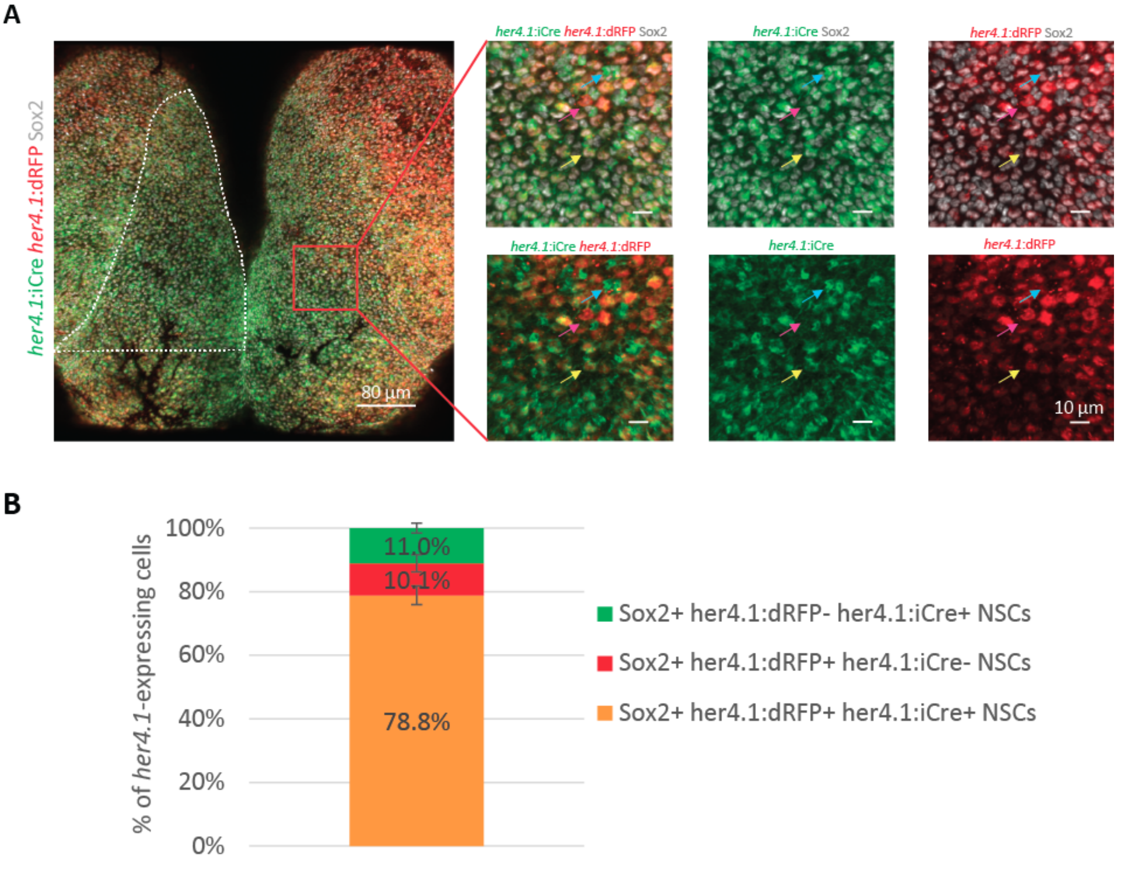


Figure S2

**Fig. S2. *her4.1*:iCre recapitulates *her4.1*:dRFP expression.** (**A**) Whole-mount dorsal view of the pallium of a *her4.1:iCre;her4.1:dRFP* double transgenic adult immunostained for iCre, dRFP and Sox2. The Dm domain analyzed is delineated by the dotted lines. Close-ups of the boxed area (split channels) illustrate dRFP$^+$/iCre$^+$ (yellow arrow), dRFP$^+$/iCre$^-$ (pink arrow) and dRFP$^-$/iCre$^+$ (blue arrow) NSCs. (**B**) Distribution of *her4.1*:dRFP$^+$ and *her4.1*:iCre$^+$ among all Sox2$^+$ cells expressing *her4.1*. n=4 brains. Error bar: s.e.m.

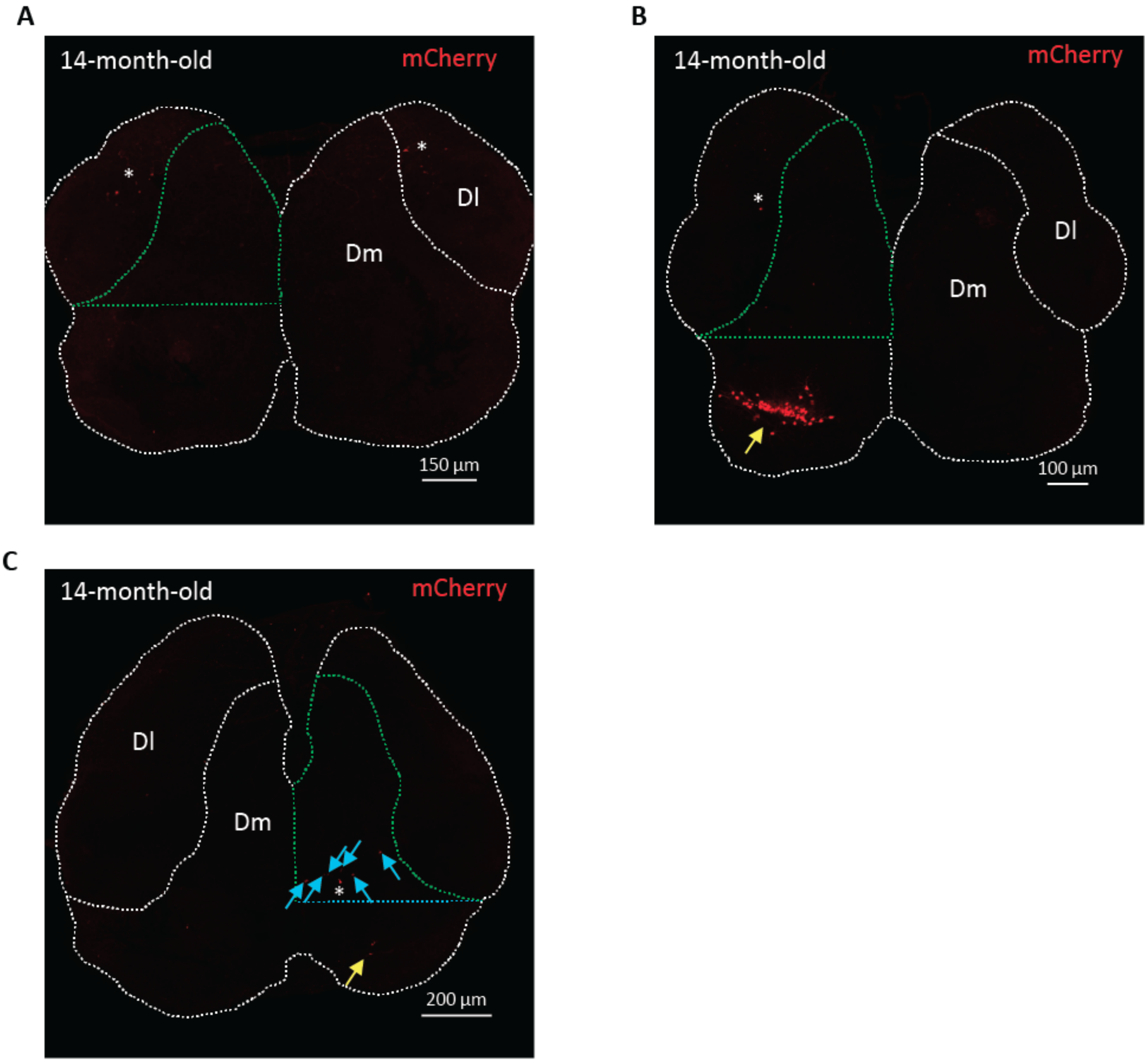


Figure S3

**Fig. S3. Absence of mCherry leakiness in the analyzed Dm region of non-induced *her4.1:iCre;ubi:Switch* zebrafish.** (**A**) Representative dorsal view of 7 out of 10 pallia showing the complete absence of recombination of the *ubi:Switch* transgene in the entire pallial area of *her4.1:iCre;ubi:Switch* uninduced double transgenic zebrafish (**B**) Dorsal view of an uninduced pallium harboring mCherry-labelled cells in its most posterior part (yellow arrow), indicating spontaneous recombination of the *ubi:Switch*. 3 out of 10 pallia displayed recombination out of the analyzed region. The pallium shown here is the one exhibiting the highest number of mCherry$^{+}$ cells. (**C**) Dorsal view of the unique case showing recombined cells in the region of interest (6 cells in a single hemisphere - blue arrows). The yellow arrow points to recombined cells located outside the region of interest. (**A** - **C**) The pallium as well as the boundary between Dm and Dl are highlighted with white dotted lines. The region analyzed in the clonal analysis is delineate in green. * mark artifacts. Fish were analyzed at one year and two months of age so as to cover most of the time span by the clonal analysis.

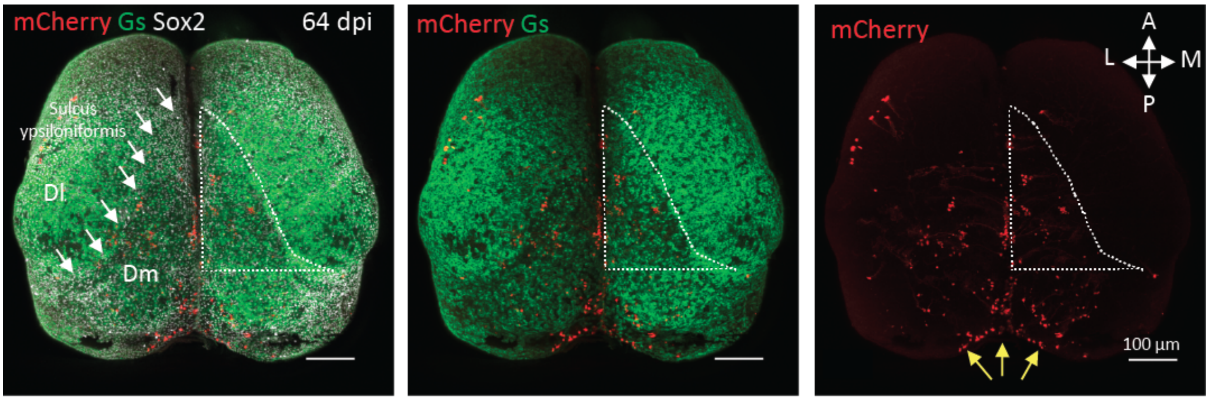


Figure S4

**Fig. S4. Delimitation of the dorso-medial pallium (Dm) region analyzed.** Dorsal whole-mount view of a pallium induced at 3 mpf and immunostained at 64 dpi for mCherry (red), Gs (green) and Sox2 (grey). White arrows show the sulcus ypsiloniformis (sy) that separates Dm from Dl. Yellow arrows point to several traced cells in the most posterior part the pallium whose clonal affiliation cannot be unambiguously ascertained. Such equivocal cell clusters resulted from a higher induction rate in this particular region and led us to exclude it from of our clonal analysis. The dotted area indicates the Dm territory selected for analysis. Its posterior border is identified by anatomical landmarks (posterior limit reached by the sy).

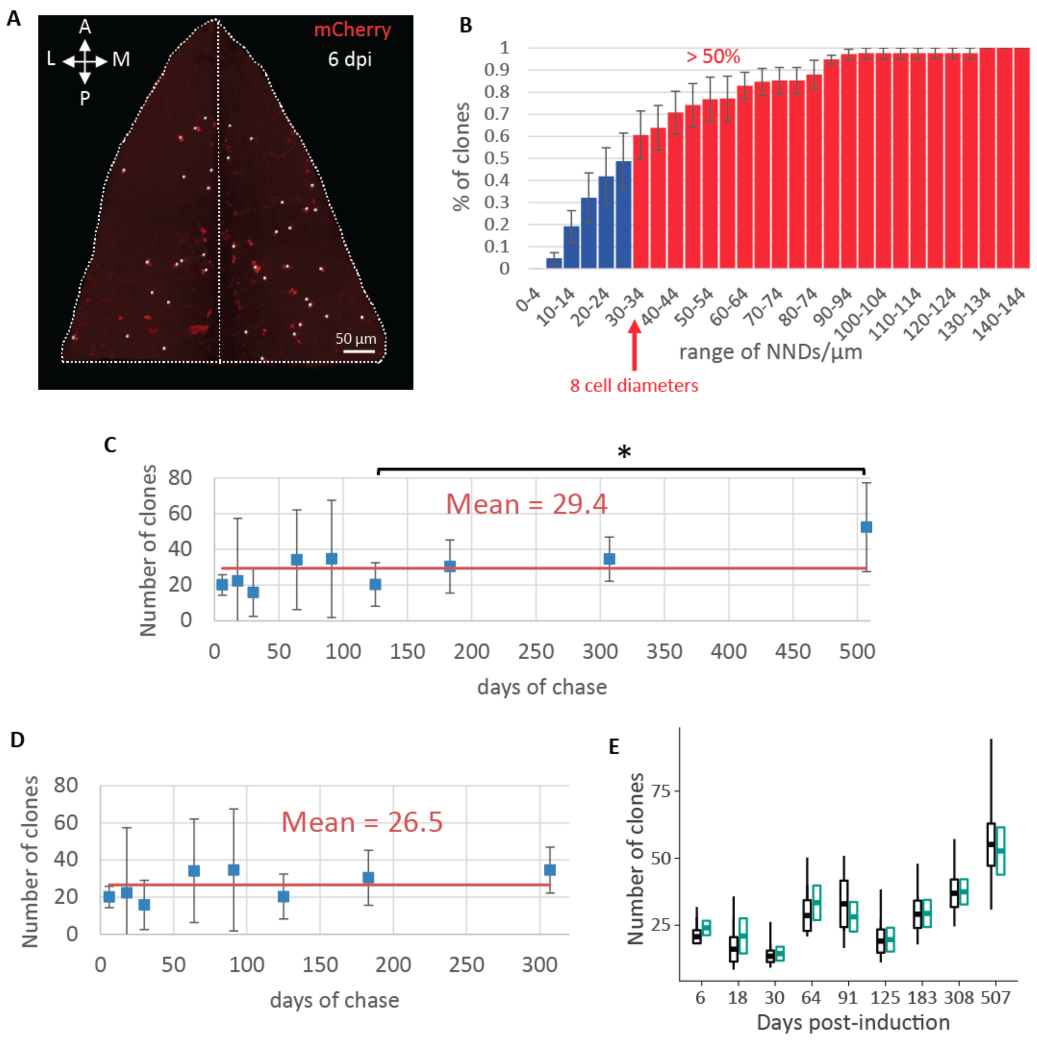


Figure S5

**Fig. S5. Clonality assessment.** (**A**) Dorsal view of the pallium at 6 dpi; mCherry$^+$ cell clusters comprising at least one Sox2$^+$ cells are highlighted with a white spot. (**B**) Cumulative probability distribution of the nearest neighbour distances (NNDs) between the Sox2$^+$ cell containing clones at 6 dpi. n=6 brains. (**C**) Average number of visually determined clones. One-way ANOVA: $F_{(8,37)}=2.76$, p=0.017; All pairwise comparisons: LSD test followed by Holm's adjustment;* p<0.05. Error bars: 95% CI. n= 6, 3, 3, 3, 5, 6, 7, 8 and 6 brains at 6, 18, 30, 64, 91, 125, 183, 307 and 507 dpi, respectively. **(D)** Same as (C) without the last time point. One-way ANOVA: $F_{(7,32)}=1.66$, p=0.15. All pairwise comparisons: LSD test followed by Holm's adjustment; p>0.05 for all comparisons. n= 6, 3, 3, 3, 5, 6, 7 and 8 brains at 6, 18, 30, 64, 91, 125, 183 and 307, respectively. Error bars: 95% CI. (**C-D**) Note the steadiness of the average number of visually determined clones after removing the last time point (507 dpi). (**E**) Number of clones determined by the clustering algorithm (black box and whisker plots). Box and whisker plots: the central bold bar and the upper and lower edges of the boxes represent respectively the mean and s.e.m. of the most likely clonal composition; the whiskers of the box correspond respectively to the 95% CI of the smallest and biggest clustering that are still within the 95% CI of the most likely clustering (see supplementary text). They reflect the combined uncertainty stemming from the clonal reconstruction and the finite sample size. The mean ± s.e.m. of the visually determined number of clones is also given for comparison (green box). n= 6, 3, 3, 3, 5, 6, 7, 8 and 6 brains at 6, 18, 30, 64, 91, 125, 183, 307 and 507 dpi, respectively.

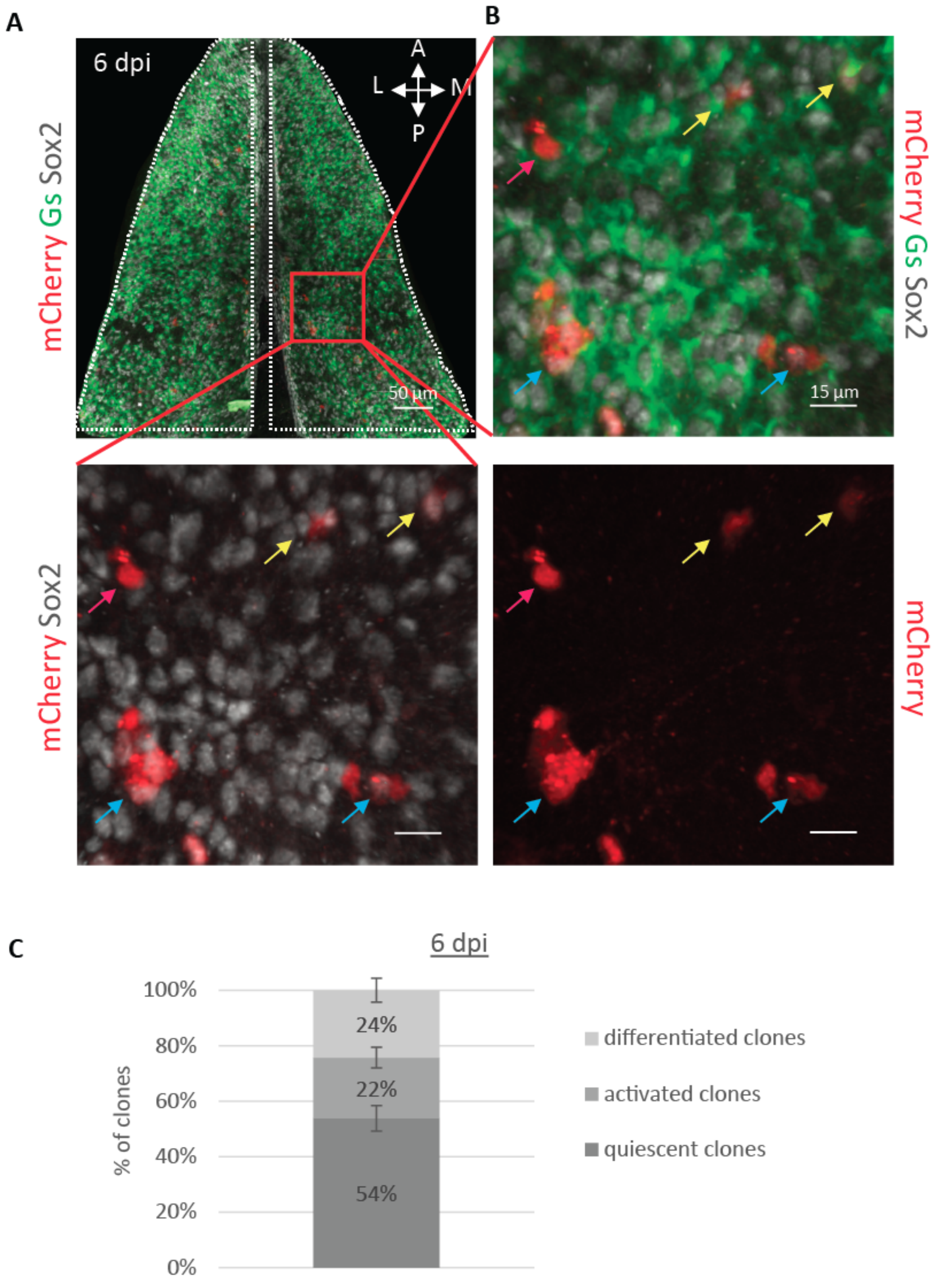


Figure S6

**Fig. S6. Activation state of clones at 6 dpi.** (**A**) Dorsal view of the pallial Dm region analyzed (seen Fig. S5) illustrating the different types of clones (red, mCherry$^{+}$) present at 6 dpi. NSCs are Gs$^{+}$ and Sox2$^{+}$, NPs are Sox2$^{+}$ only. (**B**) Higher magnification of the boxed area in (A). Blue arrows point to active clones (harboring at least two cells and comprising at least one Sox2$^{+}$ progenitor), yellow arrows to quiescent clones (comprised of a single Sox2$^{+}$ progenitor) and magenta arrows to differentiated clones (devoid of Sox2$^{+}$ progenitor). (**C**) Distribution of clones at 6 dpi based on their activation state. n=6 brains.

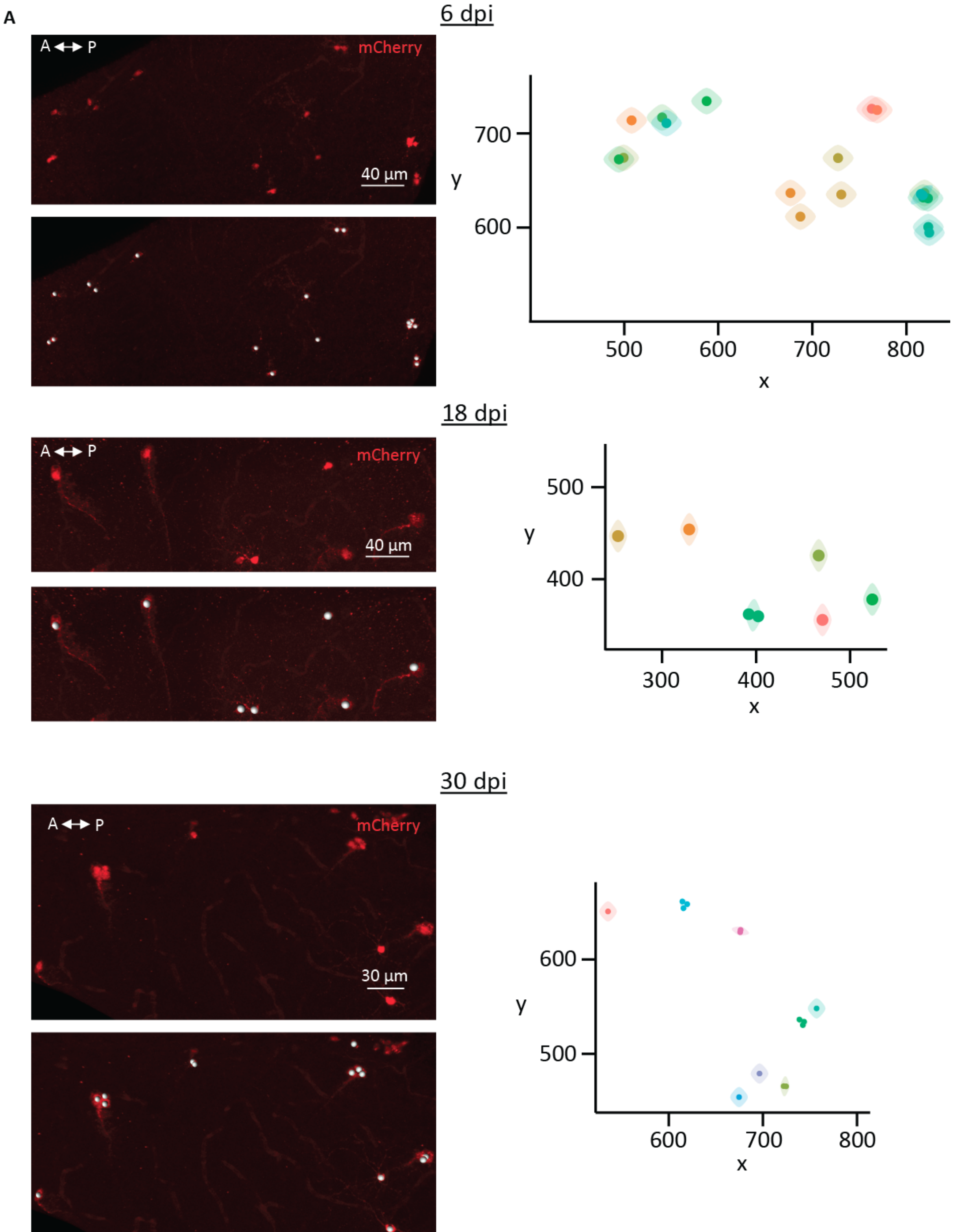

A
6 dpi
A ↔ P
mCherry
40 µm
700
600
y
500
600
700
800
x
18 dpi
A ↔ P
mCherry
40 µm
500
400
y
300
400
500
x
30 dpi
A ↔ P
mCherry
30 µm
600
500
y
600
700
800
x


Figure S7

64 dpi

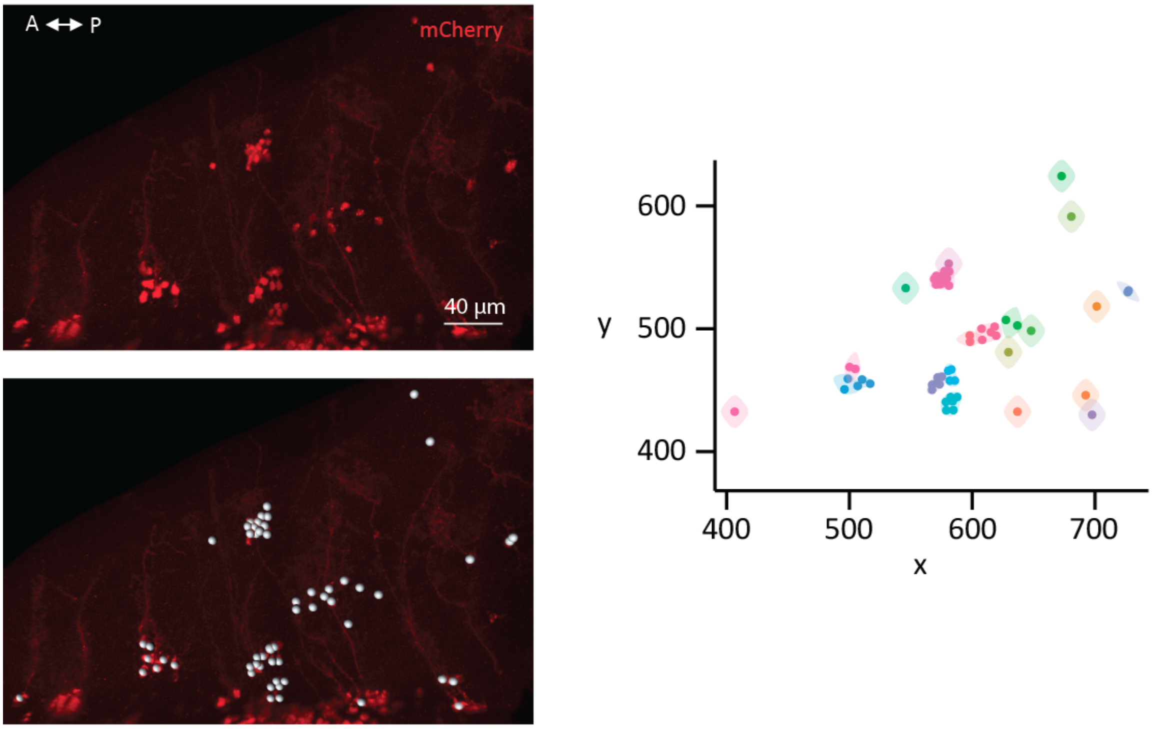


91 dpi

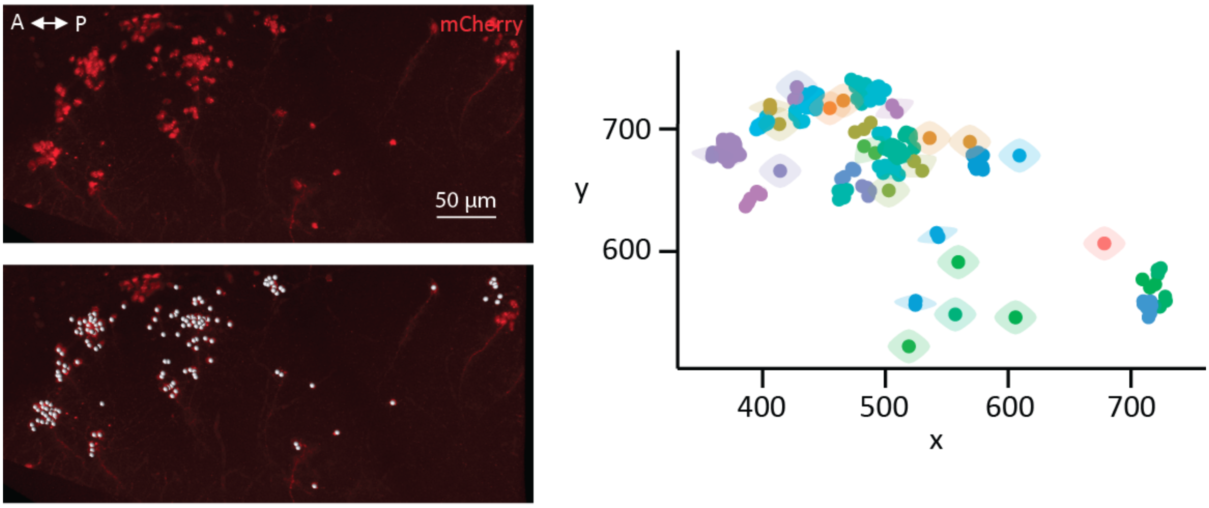


Figure S7

125 dpi

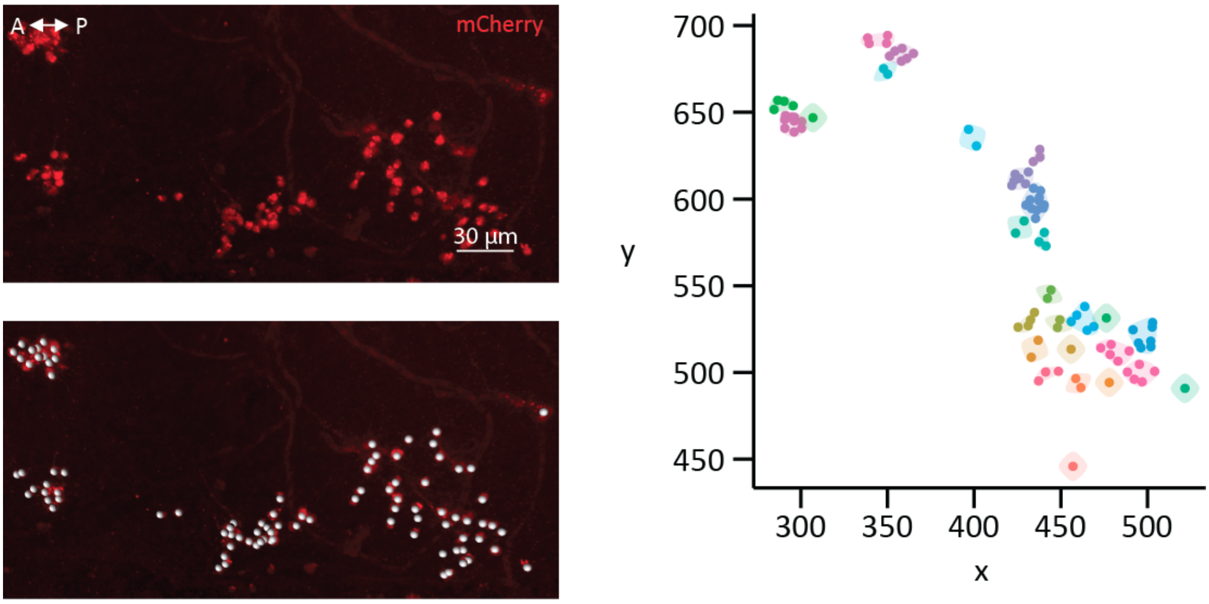


183 dpi

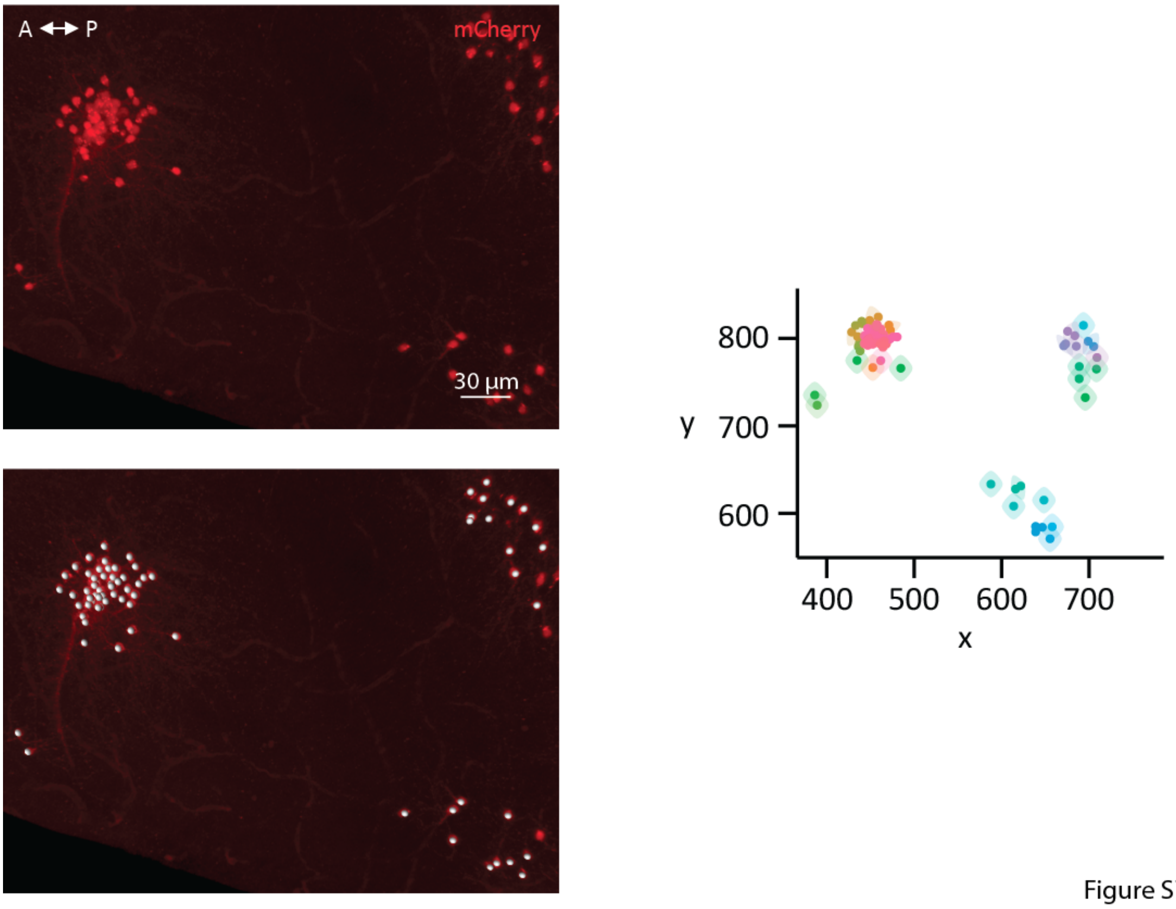


Figure S7

307 dpi

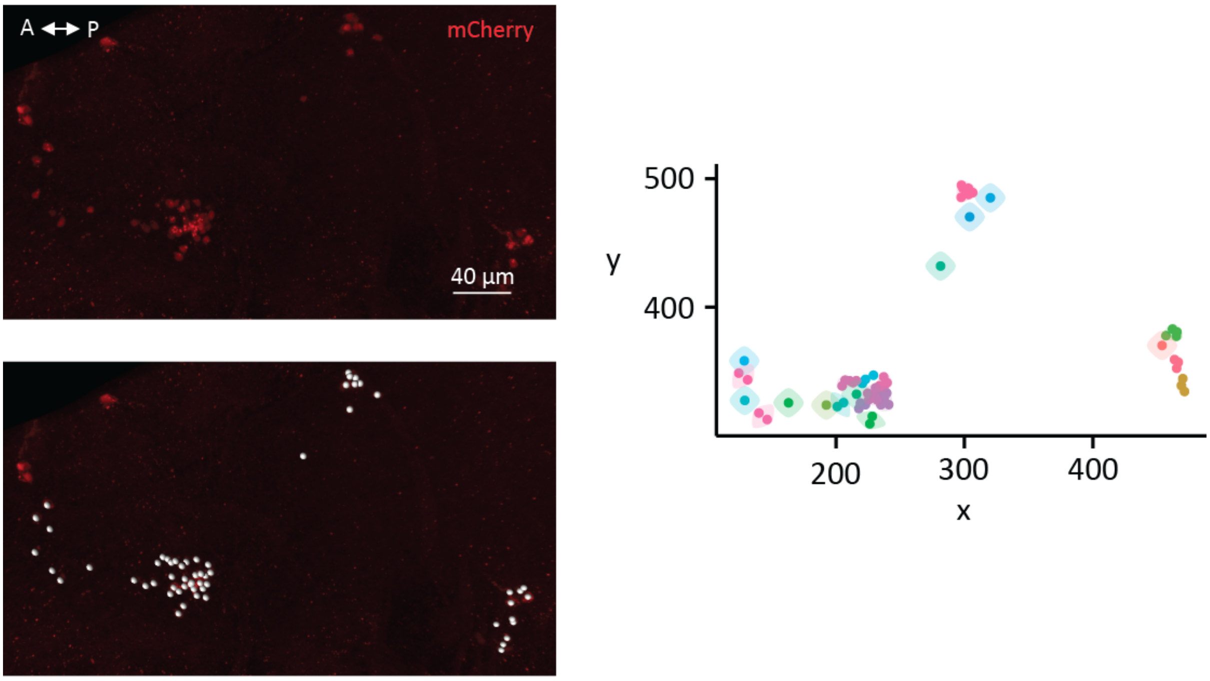


507 dpi

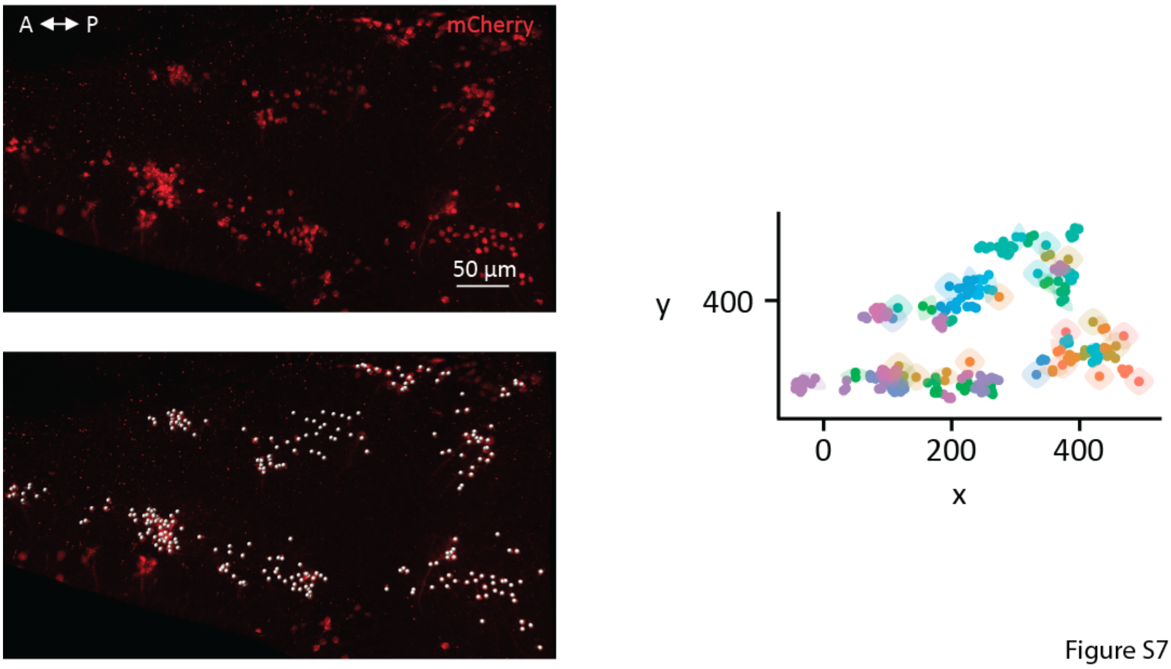


Figure S7

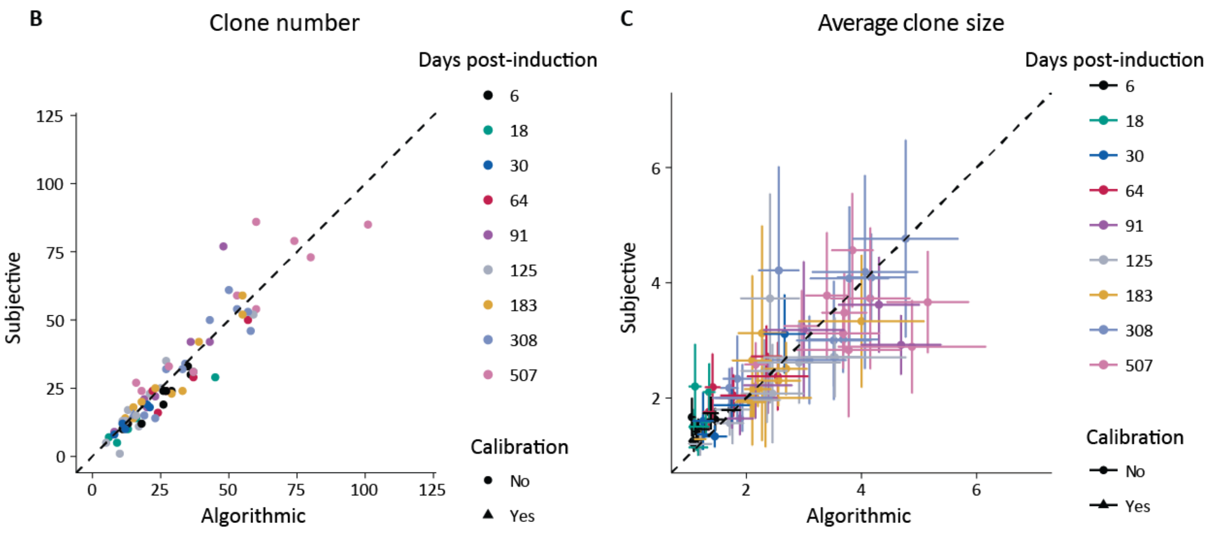


**Fig. S7. Automated cell clustering.** (**A**) Representative examples of clustering for the different time points. A dorsal view of a three-dimensional reconstruction of a hemisphere is given on the left. The mCherry channel is displayed on the upper image to show the traced cells and the spots registering their coordinates are shown superimposed on the lower image. A two-dimensional map corresponding to a planar (xy) projection of the traced cells is given on the right. Clones as determined by the clustering algorithm are color-coded. Note that a number of different clones may share the same color owing to the limited number of colors available for display. (**B**) Scatter plot showing the correlation between the numbers of clones determined by the algorithm (x-axis) and visually (y-axis). (**C**) Same plot for the average size of clones. (**B** and **C**) The different time points of the clonal analysis are represented with different colors. n= 6, 3, 3, 3, 5, 6, 7, 8 and 6 brains at 6, 18, 30, 64, 91, 125, 183, 307 and 507 dpi, respectively.

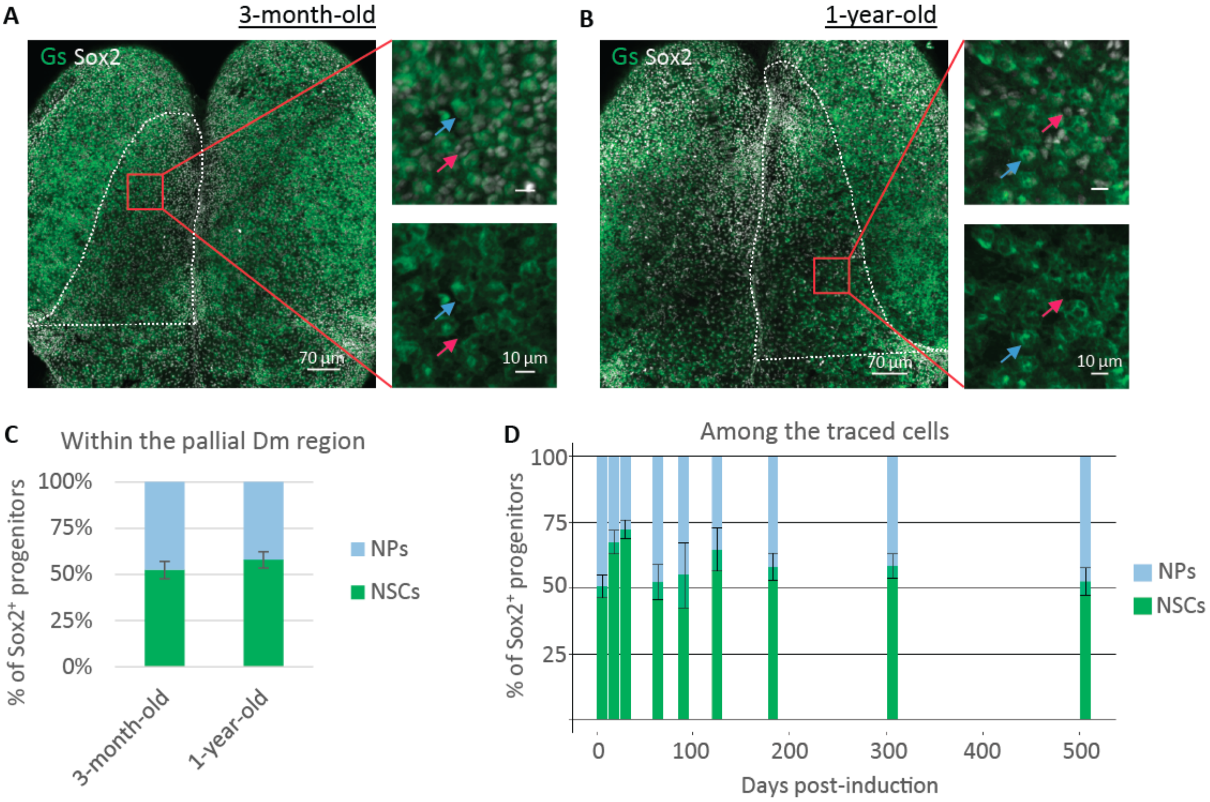


Figure S8

**Fig. S8. The behavior of Sox2$^{+}$ progenitors reads out that of NSCs.** (**A**-**B**) Dorsal view of 3 mpf (**A**) and 12 mpf (**B**) pallia immunostained for Gs and Sox2. The boxed areas are enlarged on the right to display Gs$^{+}$, Sox2$^{+}$ NSCs (blue arrow) and Gs$^{-}$, Sox2$^{+}$ NPs (pink arrow). (**C**) Quantification of the relative proportions of NSCs and NPs in the analyzed Dm region at 3 mpf and 12 mpf. p=0.43, unpaired t-test. Error bars: s.e.m. n=4 brains for both ages. (**D**) Evolution over time of the ratio of NSCs to NPs among the traced population of progenitors. One-way ANOVA: $F_{(8,37)}$=1.04, p=0.42; pairwise comparison: LSD test followed by Holm's adjustment. Error bars: s.e.m.. n= 6, 3, 3, 3, 5, 6, 7, 8 and 6 brains at 6, 18, 30, 64, 91, 125, 183, 307 and 507 dpi, respectively.

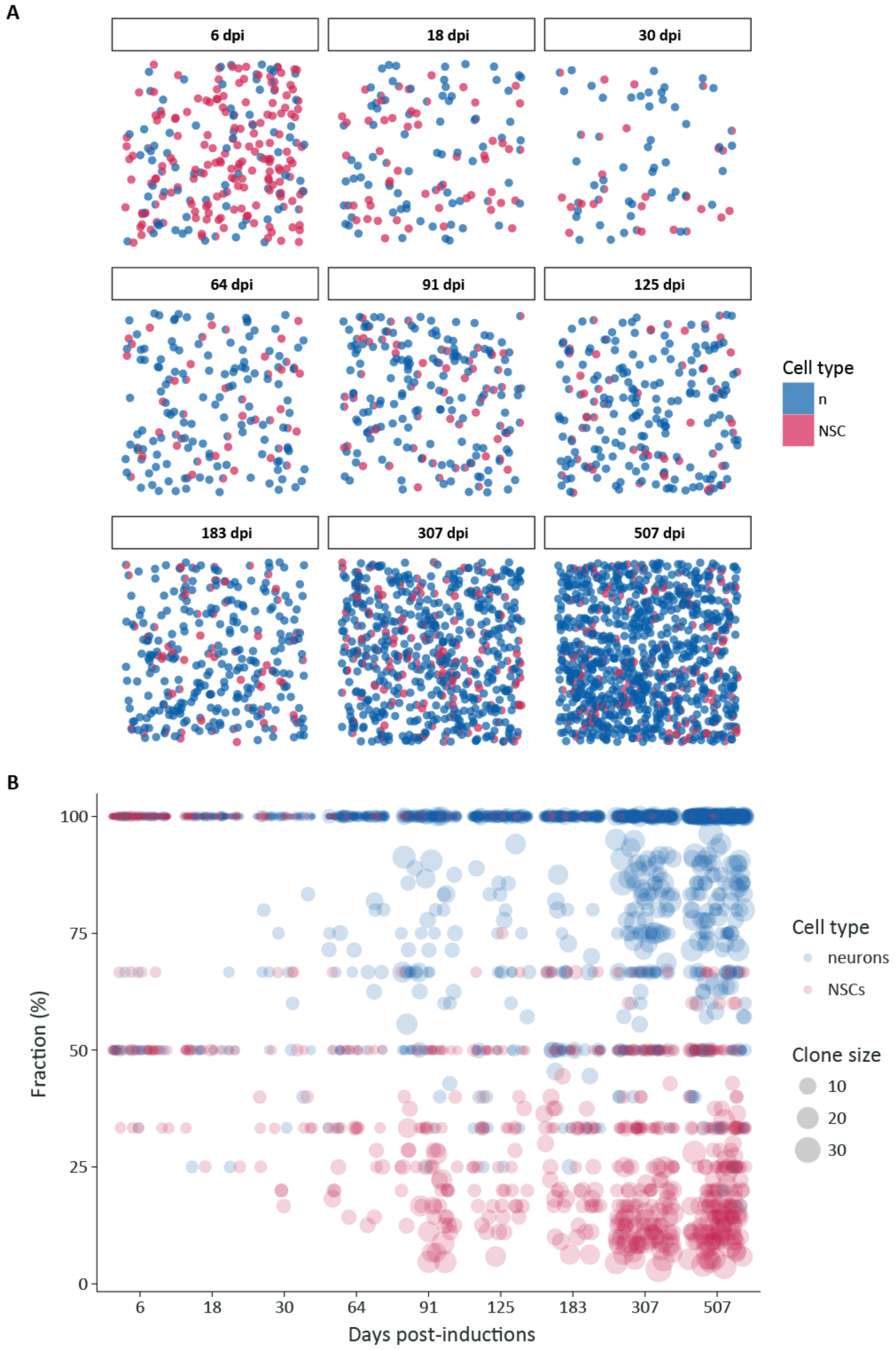


Figure S9

**Fig. S9. Clonal dynamics.** (**A**) Evolution of the clonal makeup with time. The plots display every individual clone analyzed for each time point. Each clone is represented by a pie chart showing their respective proportion of NSCs (purple) and neurons (blue). (**B**) Graphics summarizing the evolution over time of both the size and the composition of the clones. Every clone is represented by either one or two circles and every cell type in a clone has its own circle. The y-axis denotes the fraction of this cell type in the clone. Note the overall growth of the clones over time, which is concomitant with an increase in the fraction of fully differentiated clones at the expense of active clones. A small fraction of active clones expanded considerably (> 30 cells) by the end of the chase. (**A**, **B**) Clones from different fish have been pooled and their number has not been normalized. n= 6, 3, 3, 3, 5, 6, 7, 8 and 6 brains at 6, 18, 30, 64, 91, 125, 183, 307 and 507 dpi, respectively.

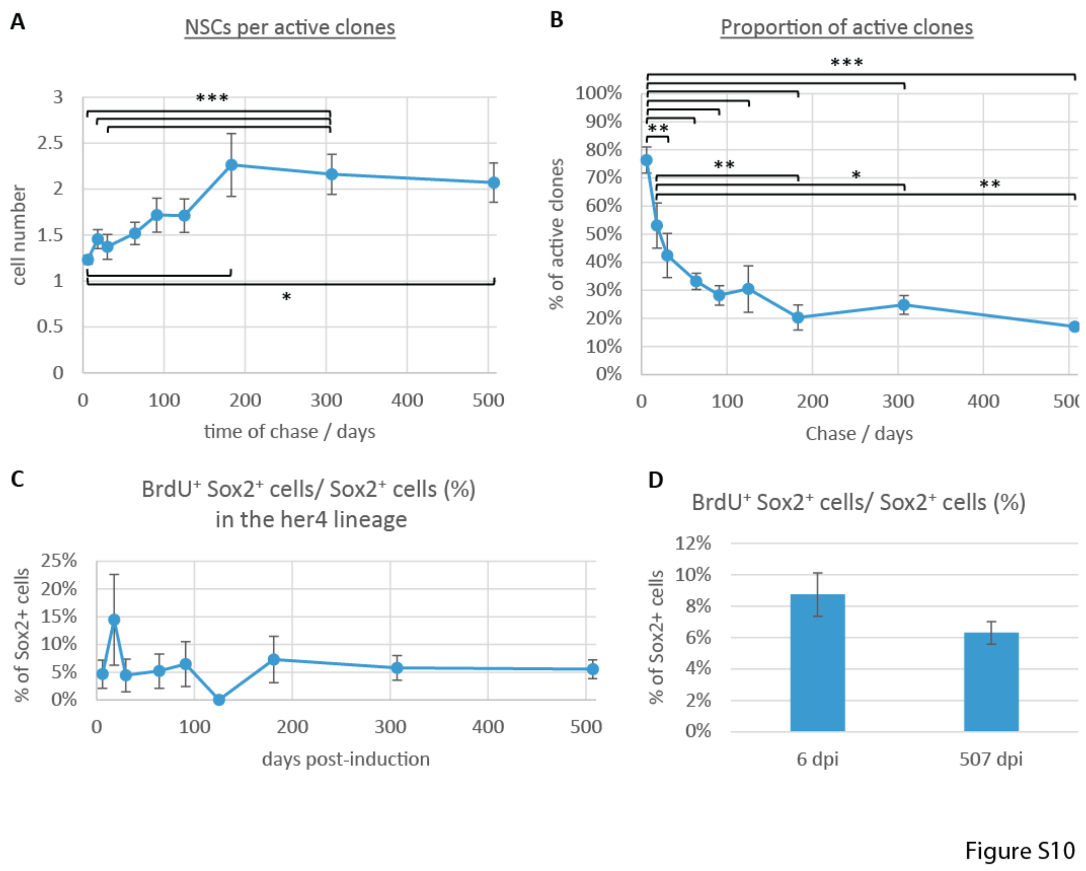


**Fig. S10. Hierarchical organization of pallial NSCs.** (**A, B**) Clonal dynamics using a visual determination of clones. (**A**) Time evolution of the number of NSCs (approximated by $Sox2^+$ progenitors) per active clone. Kruskal-Wallis test: p=0.0042; all pairwise comparisons: Behrens Fisher tests. * p< 0.05, *** p<0.001. Error bars: s.e.m. (**B**) Evolution over time of the proportion of active clones (i.e. clones maintaining at least one $Sox2^+$ progenitor). The proportion of reservoir NSCs among the traced NSCs is given by the value of the plateau (~20%). One-way ANOVA: $F_{(8,37)}=14.82$, p<0.001; all pairwise comparisons: LSD test followed by Holm's adjustment. * p< 0.05, ** p<0.01, *** p<0.001. Error bars: s.e.m. **(C)** Evolution over time of the proportion of proliferating $Sox2^+$ progenitors within the lineage. Kruskal-Wallis test: p=0.49; pairwise comparisons: p>0.05 for all comparisons (Behrens Fisher tests). Error bars: s.e.m. (**D**) Comparison of the proportion of proliferating $Sox2^+$ progenitors in the analyzed pallial area between the first and the last time point of the chase. Welch's t-test assuming unequal variances: p=0.15. (**A**-**C**) n= 6, 3, 3, 3, 5, 6, 7, 8 and 6 brains at 6, 18, 30, 64, 91, 125, 183, 307 and 507 dpi, respectively.

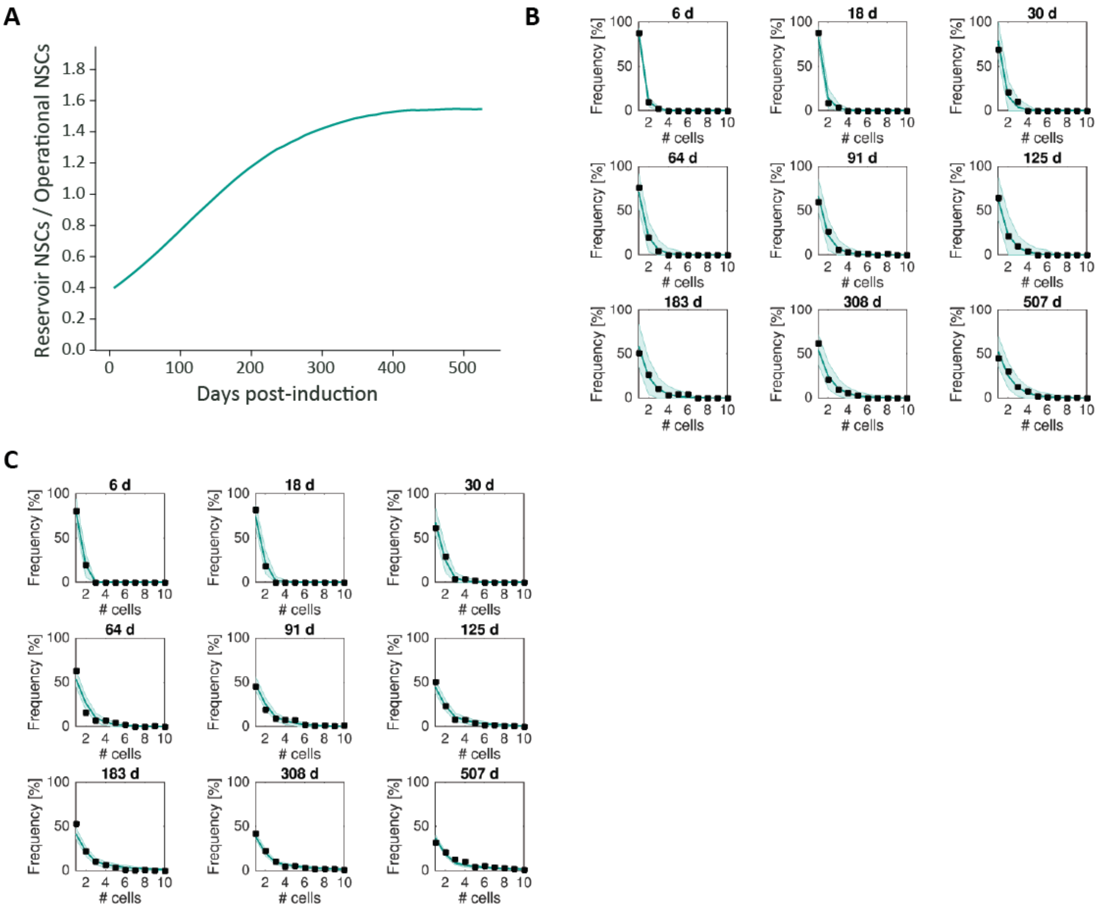


Figure S11

**Fig. S11. Clone size distributions.** (**A**) Time evolution of the ratio of reservoir to operational NSCs as inferred from the modeling. The experimental ratio at 6 dpi (0.4) was used to initialize the modeling. Because induction was biased towards operational NSCs, this ratio progressively increase from 0.4 at 6 dpi until reaching a steady state value of 1.57, which correspond to the putative actual ratio within the *her4*$^{+}$ NSC lineage. (**B**) Inverse cumulative distribution of the NSC content of the clones, indicating the chance of finding a clone with more than a given number of NSCs. (**C**) Inverse cumulative distribution of the neuronal content of the clones, indicating the chance of finding a clone with more than a given number of neurons.

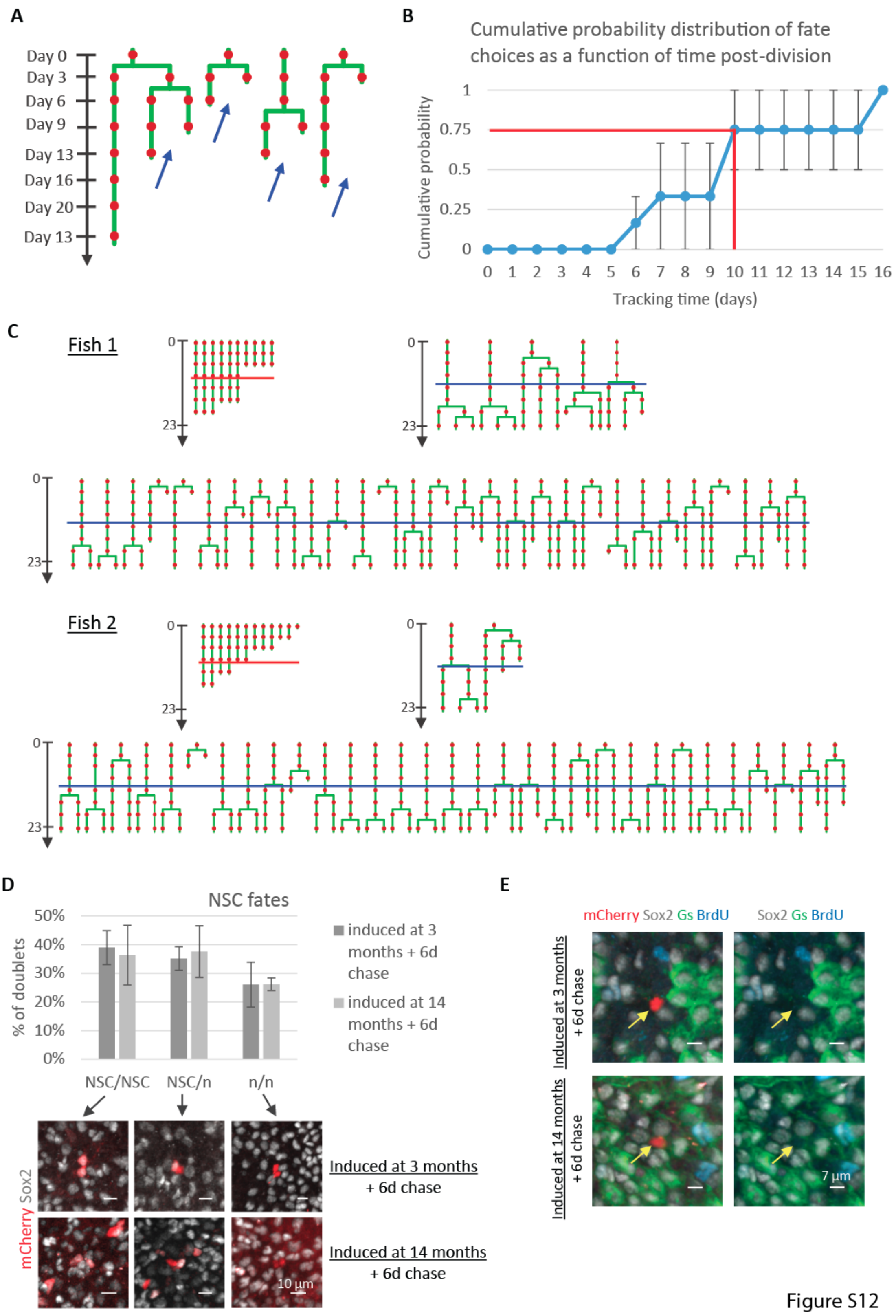


**Fig. S12. Analysis of NSC short term behavior.** (**A-B**) To determine the time necessary for the fate of a NSC to become apparent after its division, we focused on the symmetric neurogenic divisions. (**A**) Example of tracks harboring a symmetric neurogenic division (blue arrows - days of imaging sessions are indicated on the left). (**B**) Cumulative probability distribution of the time (in days) elapsed between a NSC symmetric neurogenic division and the moment when its last daughter cell lost *gfap*:dTomato expression. 75% of fate choices were apparent before ten days of tracking (red lines). Based on 3 and 2 neurogenic divisions from n=2 brains. Error bars: s.e.m.. (**C**) Illustration of all the active tracks (i.e. comprising either a division event or a direct neuronal differentiation) for two fish in the Dm territory of interest. These are a total of 44 and 45 NSCs for 322 and 373 NSCs imaged, respectively (i.e. 278 and 328 NSCs remained quiescent for the entire duration of the recording). For each fish: top left: direct differentiation tracks; top right: 2 division-tracks; bottom: 1 division-tracks. We only analyzed the outcome of divisions occurring at least three time points (i.e. about 10 days – blue bar) before the end of the experiment, and only scored direct differentiation events when taking place after at least 3 recorded time points (red bar). (**D**) Two-celled clones (doublets) recovered at 6 dpi (Fig. S4C) were used to infer NSC fate upon division ("n": neuron). The graphic shows the relative proportions of the different types of NSC divisions inferred from the doublets. Illustrations of doublets of each type are given below the graphic. Inductions were performed both in 3-month-old and 14-month-old fish and revealed no change in NSC fate with age. 3-month-old fish: n=5 brains; 14-month-old fish: n= 3. (**E**) Example of single-celled neuronal clones, expected to arise from the direct differentiation of a NSC into a neuron.

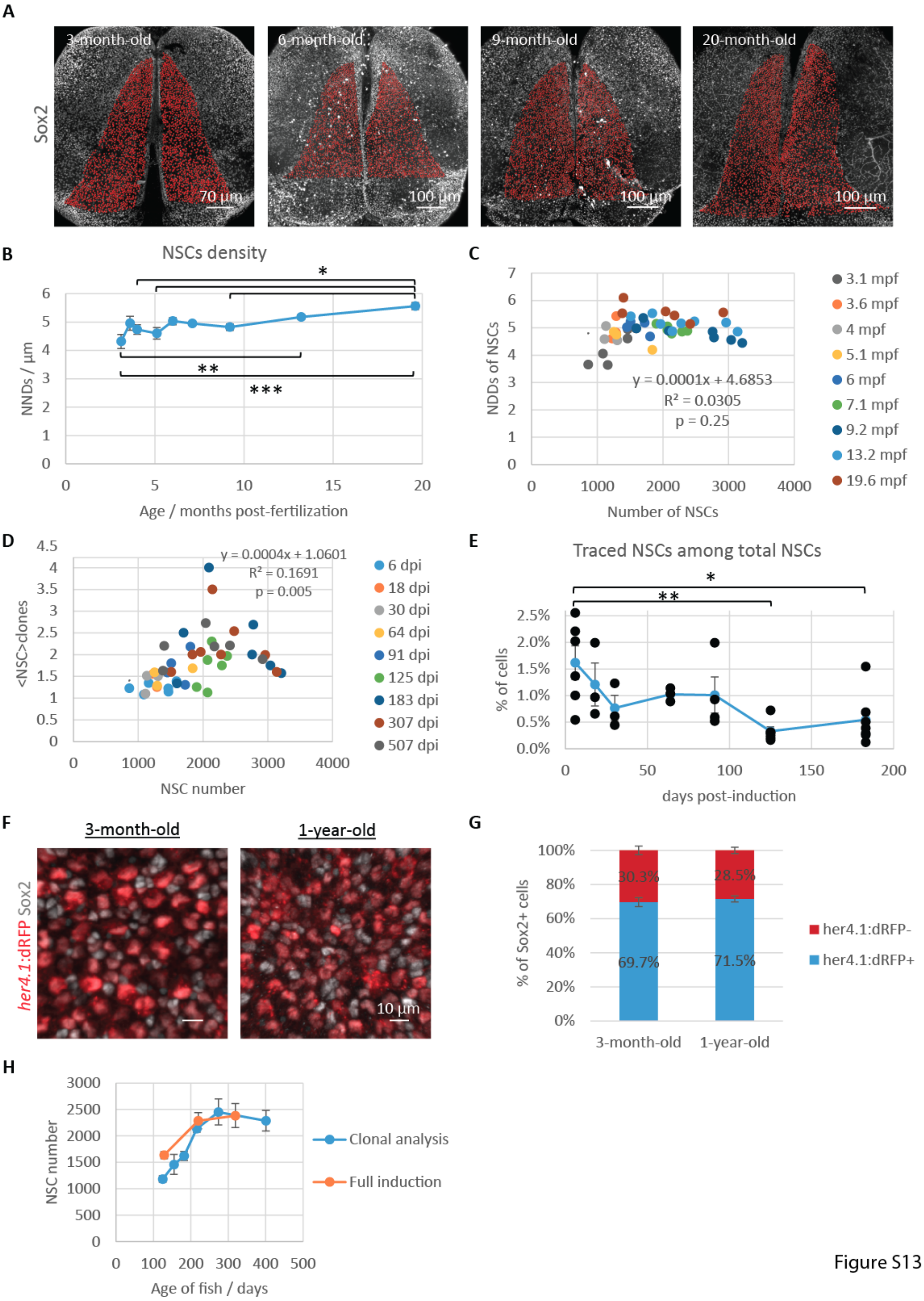


Figure S13

**Fig. S13. Ongoing production of NSCs by an upstream progenitor source.** (**A**) Dorsal view of three-dimensional reconstructions of pallia immunostained for Sox2 at different ages. All NSCs (orange dots) were counted in the Dm region of interest. (**B**) Evolution of NSC nearest neighbor distances (NNDs) with advancing age. One-way ANOVA: $F_{(8,37)}=6.09$, $p<0.001$; all pairwise comparisons: LSD test followed by Holm's adjustment. * p< 0.05, ** p<0.01, ***p<0.001. Error bars: s.e.m.. (**C**) Scatter plot showing the complete absence of correlation between the number of NSCs and their density as assessed by their NNDs. (**B**-**C**) n= 6, 3, 3, 3, 5, 6, 7, 8 and 6 brains at 3.1, 3.6, 4, 5.1, 6, 7.1, 9.2, 13.2 and 19.6 mpf, respectively. (**D**) Scatter plot presenting the relationship between the number of NSCs in the region analyzed and the NSC content of the active clones. The minor correlation (r = 0.41) between both statistics is likely fortuitous and appears insufficient to explain the expansion of the NSC population by the growth of the NSC content of the active clones. n= 6, 3, 3, 3, 5, 6, 7, 8 and 6 brains at 6, 18, 30, 64, 91, 125, 183, 307 and 507 dpi, respectively. (**E**) Evolution of the proportion of traced (mcherry$^+$) NSCs among all NSCs during their period of expansion. One-way ANOVA: $F_{(8,37)}=3.48$, $p<0.01$; all pairwise comparisons: LSD test followed by Holm's adjustment. * $p< 0.05$, ** $p<0.01$. Error bars: s.e.m. n= 6, 3, 3, 3, 5, 6 and 7 brains at 6, 18, 30, 64, 91, 125 and 183 dpi, respectively. (**F**) Dorsal views (close-ups) of the analyzed pallial region of 3-month-old and 1-year-old *her4.1*:dRFP transgenic fish immunostained for Sox2. (**G**) Respective proportion of *her4.1*:dRFP$^+$ and *her4.1*:dRFP$^-$ cells within the total Sox2$^+$ population of progenitors. Unpaired t-test: p=0.59. n=4 brains for both ages. (**H**) Comparison of the number of pallial NSCs present in fish of different ages between the clonal analysis and the full induction experiment. Error bars: s.e.m.. Clonal analysis: n= 3, 3, 5, 6, 7 and 8 brains at 125, 155, 182, 216, 274 and 401 days old, respectively. Full induction: n= 5, 6, and 5 brains at 129, 220 and 319 days old, respectively.

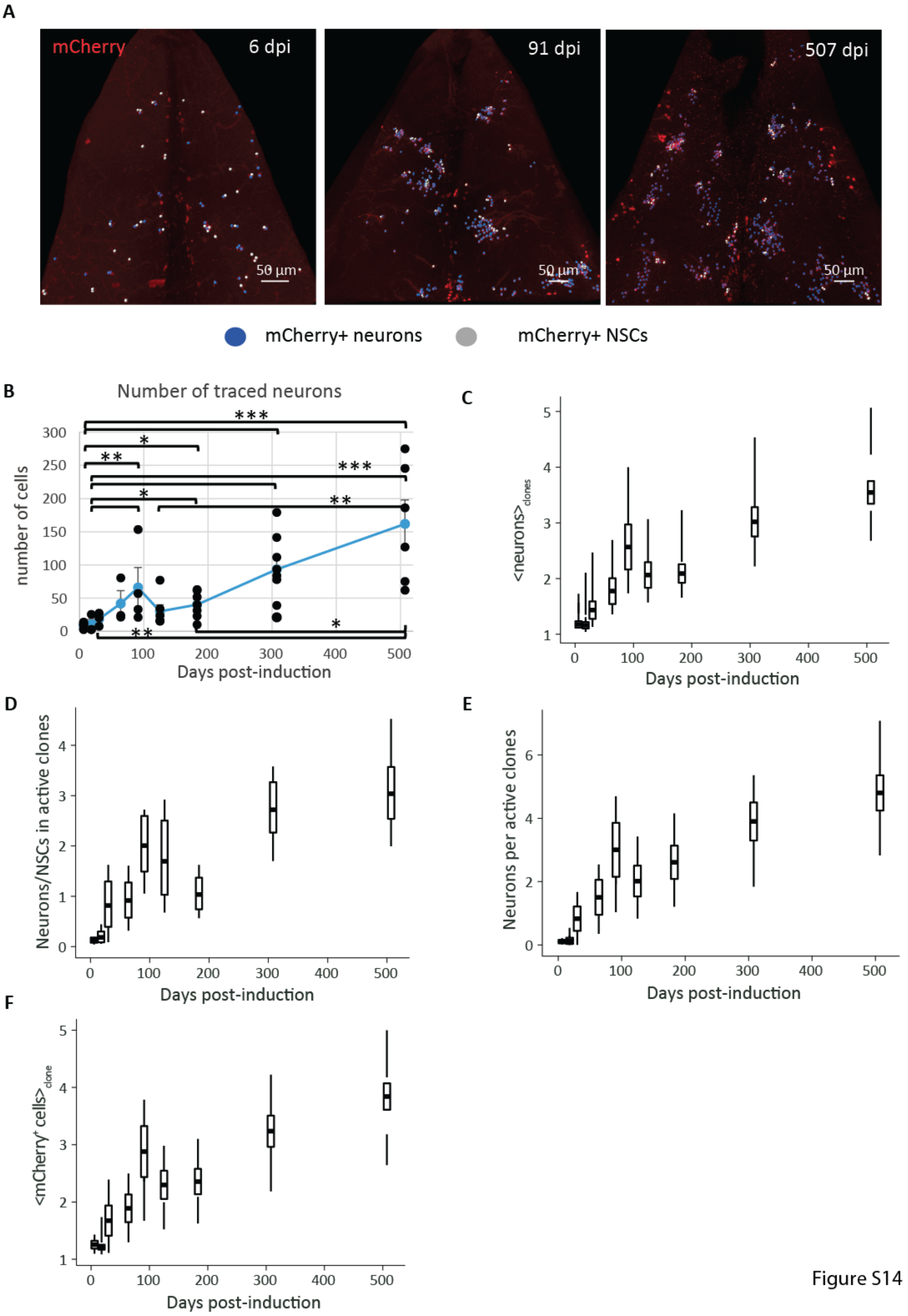


**Fig. S14. Pallial adult neurogenesis is additive in zebrafish.** (**A**) Dorsal views of the analyzed pallial Dm region at different time points of the clonal analysis. Only the mCherry channel is displayed to highlight the clones. NSCs and neurons are marked by white and blue spots, respectively. (**B**) Evolution over time of the average number of traced neurons. One-way ANOVA: $F_{(8,37)}=10.12$, $p<0.001$; all pairwise comparisons: LSD test followed by Holm's adjustment. * $p< 0.05$, ** $p<0.01$, ***$p<0.001$. Error bars: s.e.m. (**C**) Time evolution of the average number of neurons per clones. (**D**) Evolution over time of the ratio of neurons to NSCs in the active clones. (**E**) Evolution of the number of neurons per active clone across all time points analyzed. (**F**) Evolution over time of the average clone size. (**C**-**F**) Box and whisker plots: the central bold bar and the upper and lower edges of the boxes represent respectively the mean and s.e.m. of the most likely clonal composition; the whiskers of the box correspond respectively to the 95% CI of the smallest and biggest clustering that are still within the 95% CI of the most likely clustering (see supplementary text). (**B**-**G**) n= 6, 3, 3, 3, 5, 6, 7, 8 and 6 brains at 6, 18, 30, 64, 91, 125, 183, 307 and 507 dpi, respectively.

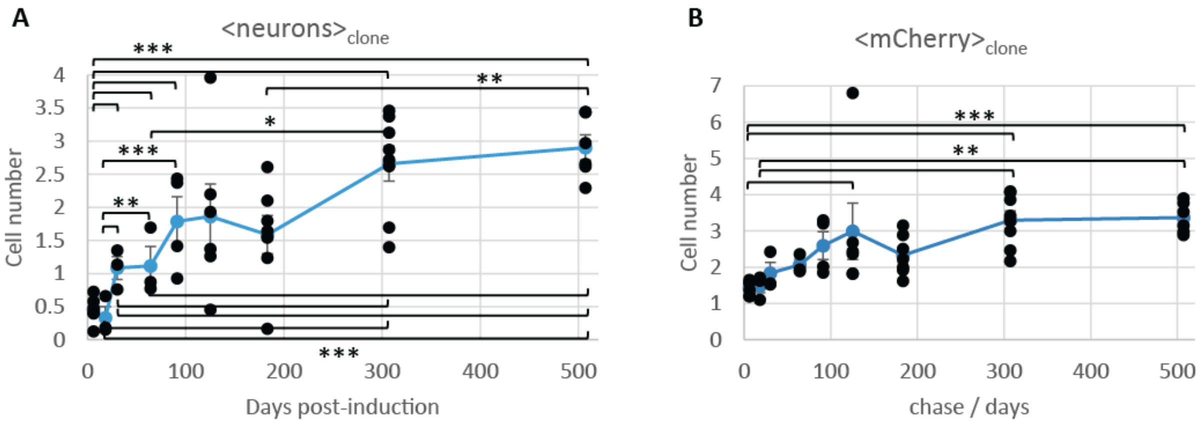


Figure S15

**Fig. S15. Pallial adult neurogenesis is additive in zebrafish.** (**A**) Evolution over time evolution of the average number of neurons per visually determined clones. Kruskal-Wallis test: $p<0.001$; all pairwise comparisons: Behrens Fisher tests. * $p< 0.05$,** $p<0.01$ and *** $p<0.001$. Error bars: s.e.m. (**B**) Evolution over time of the average clone size (visually determined clones). One-way ANOVA: $F_{(8,37)}=7.27$, $p<0.001$; all pairwise comparisons: LSD test followed by Holm's adjustment. ** $p<0.01$, ***$p<0.001$. Error bars: s.e.m. (**A**-**B**) n= 6, 3, 3, 3, 5, 6, 7, 8 and 6 brains at 6, 18, 30, 64, 91, 125, 183, 307 and 507 dpi, respectively.

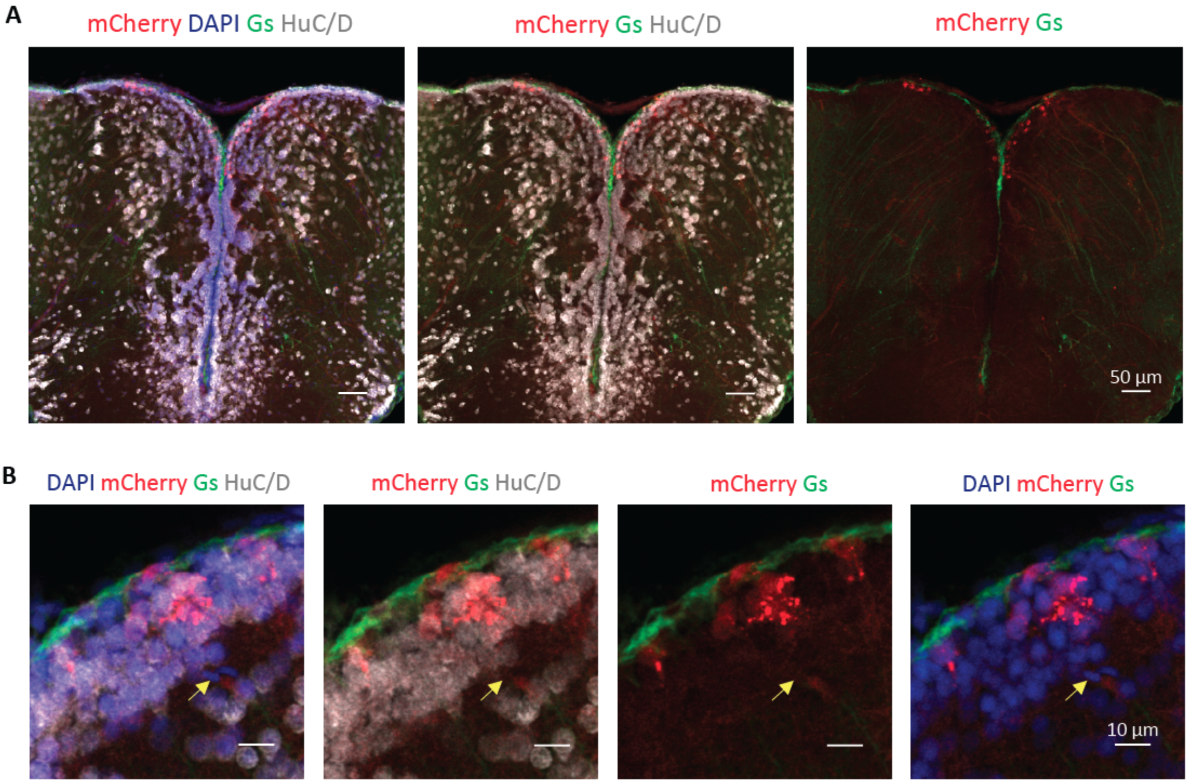


Figure S16

**Fig. S16. Parenchymal traced cells are neurons.** (**A**) Cross-section of the telencephalon of an *her4.1:iCre;ubi:Switch* fish induced at 3 months and immunostained for mCherry (red), Gs (green) and neuronal marker HuC/D (grey). Note that almost all the parenchymal cells stain for HuC/D. (**B**) Closeup on some traced cells. The yellow arrow points to putative endothelial cells recognizable by their small flattened nuclei.

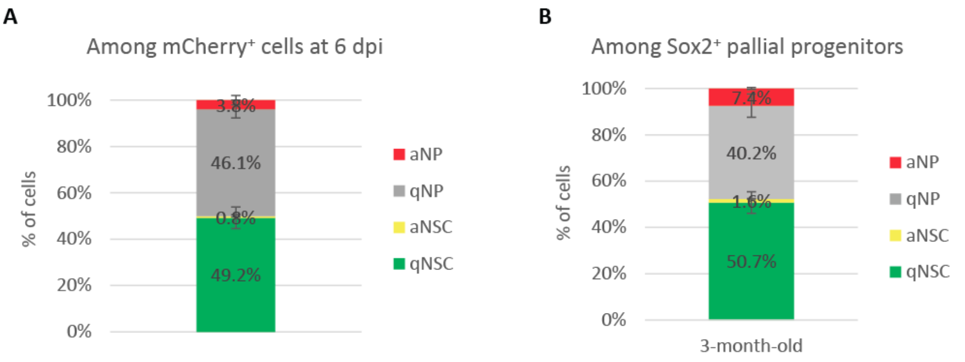


Figure S17

**Fig. S17. Characterization of the traced population of progenitors.** (**A**) Progenitor composition of the mCherry$^+$ population at 6 dpi. n=6 brains. (**B**) Distribution of the different types of progenitors within the pallial Dm region. n=4 brains.

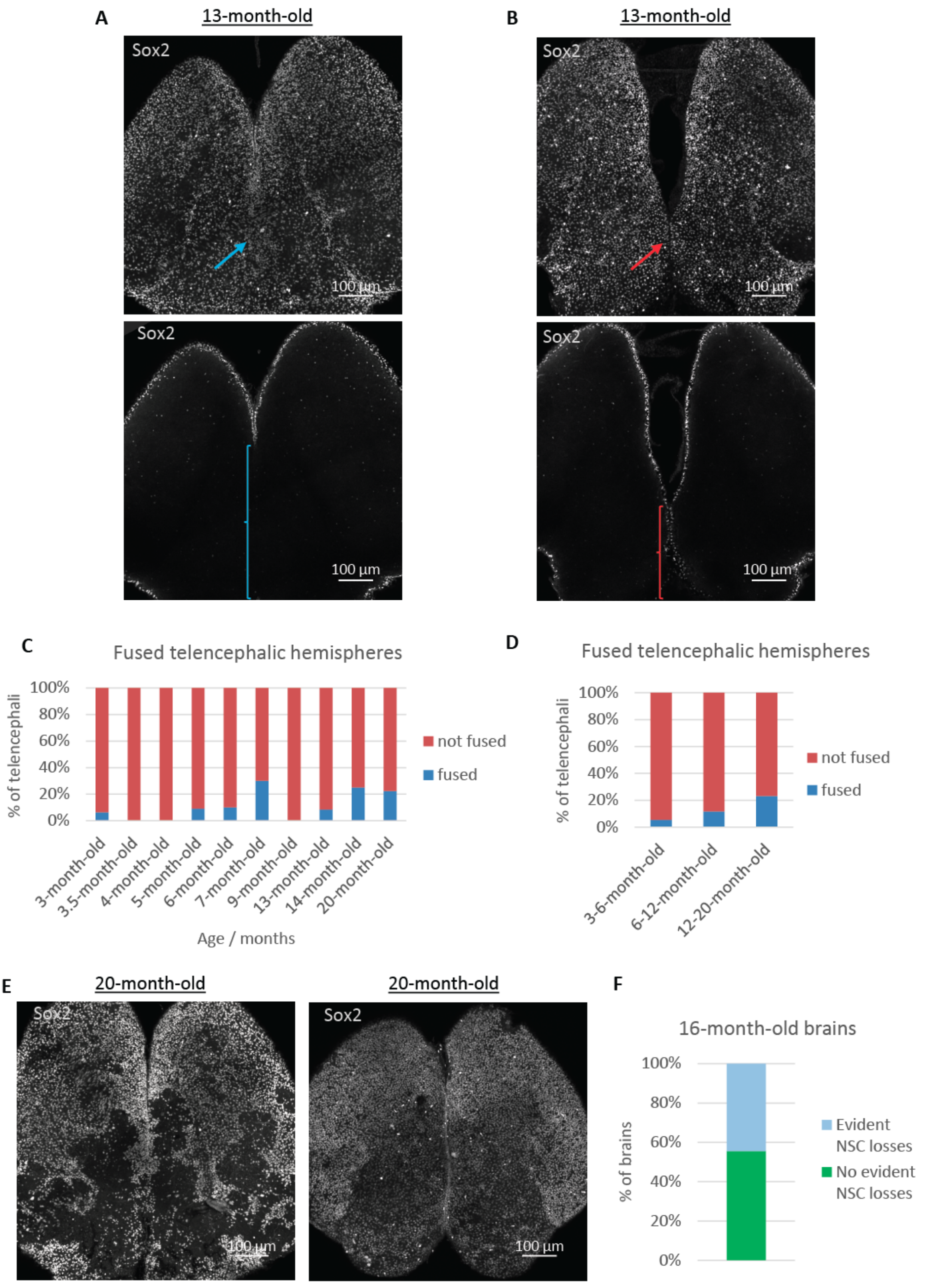


Figure S18

**Fig. S18. The aging pallium.** (**A**) Top: dorsal view of a three-dimensional reconstruction of a pallium from a 13-month-old fish displaying an interhemispheric fusion (blue arrow). Sox2 immunostaining is shown in grey. Bottom: optical horizontal section showing the region of the fusion (blue brace). (**B**) Same as in (A) for a pallium that does not harbor any interhemispheric fusion. The red arrow and brace highlight the junction between both hemispheres on the three-dimensional reconstruction and on the optical section of the pallium, respectively. (**C**) Quantification of the proportion of brains displaying interhemispheric fusions across all ages analyzed in the clonal analysis. Fisher's exact test: p=0.35. n=16, 8, 10, 11, 10, 10, 9, 24 and 9 for 3-, 3.5-, 4-, 5-, 6-, 7-, 9-, 13-, 14- and 20-month-old fish, respectively. (**D**) Quantification of the proportion of brains of different age categories displaying interhemispheric fusions. Fisher's exact test: p=0.11. n=55, 43 and 13 for 3-6, 6-12 and 12-20 months old fish, respectively. (**E**) Left: dorsal view of a three-dimensional reconstruction of a pallium from a 13-month-old fish harboring evident NSC losses. Sox2 immunostaining is shown in grey. Right: dorsal view of a three-dimensional reconstruction of a pallium from a 13-month-old fish that does not display any obvious NSC loss. (**F**) Quantification of the proportion of brains displaying evident NSC losses. n=9. (**C**-**D, F**) Note that more brains than included in the clonal analysis were assessed.
